# Neryl diphosphate-derived monoterpene biosynthesis via a biosynthetic gene cluster in the liverwort *Marchantia polymorpha*

**DOI:** 10.64898/2026.07.16.738752

**Authors:** Guo Wei, Tomoko Kawaguchi, Facundo Romani, Eduardo Flores-Sandoval, Meng Xie, Xinlu Chen, Takao Koeduka, Kan-ichi Iwasaki, Mutsumi Nakanishi, Hisashi Hemmi, Jin-Gui Chen, John L. Bowman, Jim Haseloff, Kenji Matsui, Feng Chen

## Abstract

Monoterpenes (C_10_) are a large group of specialized metabolites important for plant interactions with the environment. Their biosynthesis is well understood in seed plants, where geranyl diphosphate serves as the canonical substrate, but knowledge of monoterpene biosynthesis outside seed plants remains very limited. Here, we report neryl diphosphate (NPP)-derived monoterpene biosynthesis via a biosynthetic gene cluster in the liverwort *Marchantia polymorpha*. MpMTPSL2, a microbial-type terpene synthase, converts NPP into α-phellandrene and D-limonene *in vitro*. CRISPR knockout lines showed reduced production of both monoterpenes, providing direct genetic evidence for its *in planta* function. *MpCPT5*, a cis-prenyltransferase (CPT) family member identified through co-expression with *MpMTPSL2*, was confirmed to encode NPP synthase, as its knockout plants abolished α-phellandrene and D-limonene production. Subcellular localization analyses in protoplasts and stable transgenic plants demonstrated that both MpCPT5 and MpMTPSL2 localize to plastids, co-localizing across all cell types with markedly stronger signals in non-green plastids of oil-body cells. Consistent with this, expression of both genes under their respective promoters was nearly abolished in oil-body-deficient mutants and strongly upregulated in a gain-of-function line for oil-body formation. *MpMTPSL2* and *MpCPT5* are physically linked through a shared bidirectional promoter that drives their coexpression specific to oil body cells, forming a unique biosynthetic gene cluster whose coordinated expression is maintained by PRC2-mediated H3K27me3 repression. Phylogenetic analysis implies that NPP synthases in *M. polymorpha* and in flowering plants evolved independently from their respective long-chain CPT ancestors. These findings provide new insights into the mechanisms and evolution of monoterpene biosynthesis in non-seed plants.

**Significance statement:** Monoterpenes are a diverse group of specialized metabolites produced widely among land plants, yet our understanding of their biosynthesis outside seed plants remains limited. Here we report that in the liverwort *Marchantia polymorpha*, the non-canonical substrate neryl diphosphate is used for monoterpene biosynthesis. The functions of the monoterpene synthase gene *MpMTPSL2* and the neryl diphosphate synthase gene *MpCPT5* were demonstrated through CRISPR knockouts. These two genes are physically linked and share a bidirectional promoter. Promoter assays show both genes function in plastids within oil-body cells, revealing cell-type specificity. These findings shew new light on the mechanisms and evolution of monoterpene biosynthesis in non-seed plants.

## Introduction

Monoterpenes are a major subclass of terpenoids, which constitute the largest family of secondary metabolites made by land plants (1). Monoterpenes function as key chemical signals mediating interactions between plants and their environment (2), playing roles in defenses against natural enemies and attraction of beneficial organisms (3). Biosynthesis of monoterpenes has been extensively studied in seed plants, where typical plant terpene synthases (TPSs) are pivotal enzymes. TPSs form a mid-sized family in land plants and can be further divided into a, b, c, d, e/f, g and h subfamilies (1). In gymnosperms, monoterpene biosynthesis is catalyzed by members of d1 group of the TPS-d subfamily (4). In angiosperms, monoterpenes are synthesized mainly by members of the TPS-b and TPS-g subfamilies (1). Geranyl diphosphate (GPP) is the canonical substrate for monoterpenes in seed plants. In a few flowering plants, neryl diphosphate, the *cis*-isomer of GPP, was demonstrated to serve as an additional substate for monoterpenes (5). In contrast, our understanding of monoterpene biosynthesis outside seed plants is very limited. Archegoniate plants, comprising all land plant lineages except seed plants, contain both *TPS* genes and microbial terpene synthase-like (*MTPSL*) genes (6–8). TPSs of archegoniate plants have been demonstrated to encode diterpene synthase (6, 8). As such, monoterpene biosynthesis in archegoniate plants has been hypothesized to be catalyzed by MTPSLs (8). In a few archegoniate plants, several MTPSLs have been shown to have monoterpene synthase activities in *in vitro* enzyme assays using either GPP or NPP as substrate (6, 8, 9). However, genetic evidence for the role of MTPSLs in monoterpene biosynthesis in archegoniate plants is largely lacking.

GPP and NPP are synthesized by enzymes of distinct protein families. The enzyme for synthesizing GPP, GPP synthase (GPPS), belongs to the *trans*-isoprenyl diphosphate synthase (IDS) family. There are two distinct types of GPPS: homomeric GPPS and heteromeric GPPS, both of which use dimethylallyl diphosphate (DMAPP) and isopentenyl diphosphate (IPP) from the 2-C-methyl-D-erythritol 4-phosphate (MEP) pathway as substrates to yield GPP (10, 11). Heteromeric GPPSs consist of large (GPPS.LSU) and small subunits (GPPS.SSU) (12–14). GPPS is proposed to have evolved from geranylgeranyl diphosphate synthase (GGPPS) with the latter synthesizing GGPP (15). In contrast, NPP is synthesized by NPP synthase (NPPS), which belongs to the *cis*-prenyltransferase (CPT) family (16). IDSs and CPTs do not share sequence homology, indicating distinct evolutionary origins (17). NPPS is hypothesized to have evolved from long-chain CPTs of essential functions, particularly for the biosynthesis of polyprenols (16). Some GGPPSs of non-seed plants have been shown to have moonlighting GPPS activities (15). This raises an intriguing question about monoterpene biosynthesis in non-seed plants: does GPP produced by the moonlighting GGPPS serve as substate for monoterpenes or does NPP serve as a substrate?

In this study we choose *Marchantia polymorpha* as a model archegoniate plant to investigate monoterpene biosynthesis. *M. polymorpha* is a liverwort, which together with mosses and hornworts form the group of bryophytes. Bryophytes share a common ancestor with seed plants over 450 million years ago (18). Among the three lineages, liverworts are well known for their ability of producing diverse terpenes (19). *M. polymorpha*, the first liverwort to have its genome sequenced (20) and known for its genetic tractability (18), has served as a model for studying terpene biosynthesis. Like many other liverworts, *M. polymorpha* produces diverse terpenes (9). *In vitro* biochemical evidence indicates that *M. polymorpha TPS* genes encode diterpenes synthases (9, 21) while the enzymes encoded by *MTPSL* genes have sesquiterpene or monoterpene synthase activities (7–9). In this study, we sought to establish direct genetic evidence for the role of MTPSL enzymes in monoterpene biosynthesis using transgenic approaches. More importantly, we aim to identify the gene for the production of the substrate for monoterpenes in *M. polymorpha* and to understand its evolution.

A hallmark of liverwort biology is the presence of oil bodies, which are membrane-bound, specialized organelles unique to this lineage that serve as the primary storage compartments for an array of lipophilic specialized metabolites, including terpenoids (22). Genetic studies in *M. polymorpha* have shown that loss-of-function mutants lacking oil bodies exhibit dramatically reduced accumulation of terpenoids, establishing a direct link between oil body integrity and terpenoid storage (23). Furthermore, several sesquiterpene biosynthetic enzymes have been localized to oil body–containing cells, suggesting that these organelles may function not merely as passive reservoirs but as active sites of terpenoid biosynthesis (24). Despite these advances, the relationship between oil bodies and monoterpene biosynthesis remains unresolved. Therefore, after the identification of monoterpene synthase and the gene for producing its substrate, it is also our objective in this study to determine whether monoterpene biosynthesis occurs in oil body cells. All the findings in *M. polymorpha* will provide new insights into the mechanisms and evolution of monoterpene biosynthesis in land plants.

## RESULTS

### Production and analysis of Mp*mtpsl2* loss-of-function mutants

Among the terpene synthase genes in *M. polymorpha* that we have been previously characterized (9, 21), MpMTPSL2 was demonstrated to encode an enzyme catalyzing the formation of D-limonene from NPP in *in vitro* enzyme assays. As D-limonene was the most abundant monoterpene detected from *M. polymorpha* plants (9), this association suggests that MpMTPSL2 is one of the major genes for monoterpene biosynthesis in this plant. In our new *in vitro* enzyme assays, we discovered that in addition to D-limonene, recombinant MpMTPSL2 also produces a minor monoterpene α-phellandrene (*SI Appendix*, Fig. S1). To obtain genetic evidence for the *in planta* function of Mp*MTPSL2*, Mp*MTPSL2* loss-of-function mutants were produced using CRISPR/Cas9 using gRNAs targeting the beginning of the terpene synthase protein domain. Two independent alleles, Mp*mtpsl2-1* and Mp*mtpsl2-11*, were identified and analyzed (Fig. 1A). The Mp*mtpsl2-1* allele contains a 1 bp insertion in the first exon (*SI Appendix*, Fig. S2). The Mp*mtpsl2-11* allele contains a 1 bp deletion and a 31 bp insertion in the first exon (*SI Appendix*, Fig. S3). As a result, the Mp*mtpsl2-1* allele and the Mp*mtpsl2-11* allele encode proteins of 134 amino acids and 454 amino acids, respectively, in contrast to the 444 amino acid protein encoded by the wild-type Mp*MTPSL2* (*SI Appendix*, Fig. S4). Under normal growing conditions, both germinating gemmae (Fig. 1B) and mature mutant plants (Fig. 1C) are morphologically similar to the wild-type Tak-1 plants. In germinating gemmae of wild-type Tak-1 plants, α-phellandrene and D-limonene concentrations peaked at 6 days after germination and then declined rapidly by 8 and 10 days after germination (*SI Appendix*, Fig. S5). In gemmae of Mp*mtpsl2* mutant lines at 6 days after germination, α-phellandrene was undetectable (Fig. 1D), while D-limonene levels were reduced by 81.3% compared with wild-type plants (Fig. 1E)

**Figure 1.**
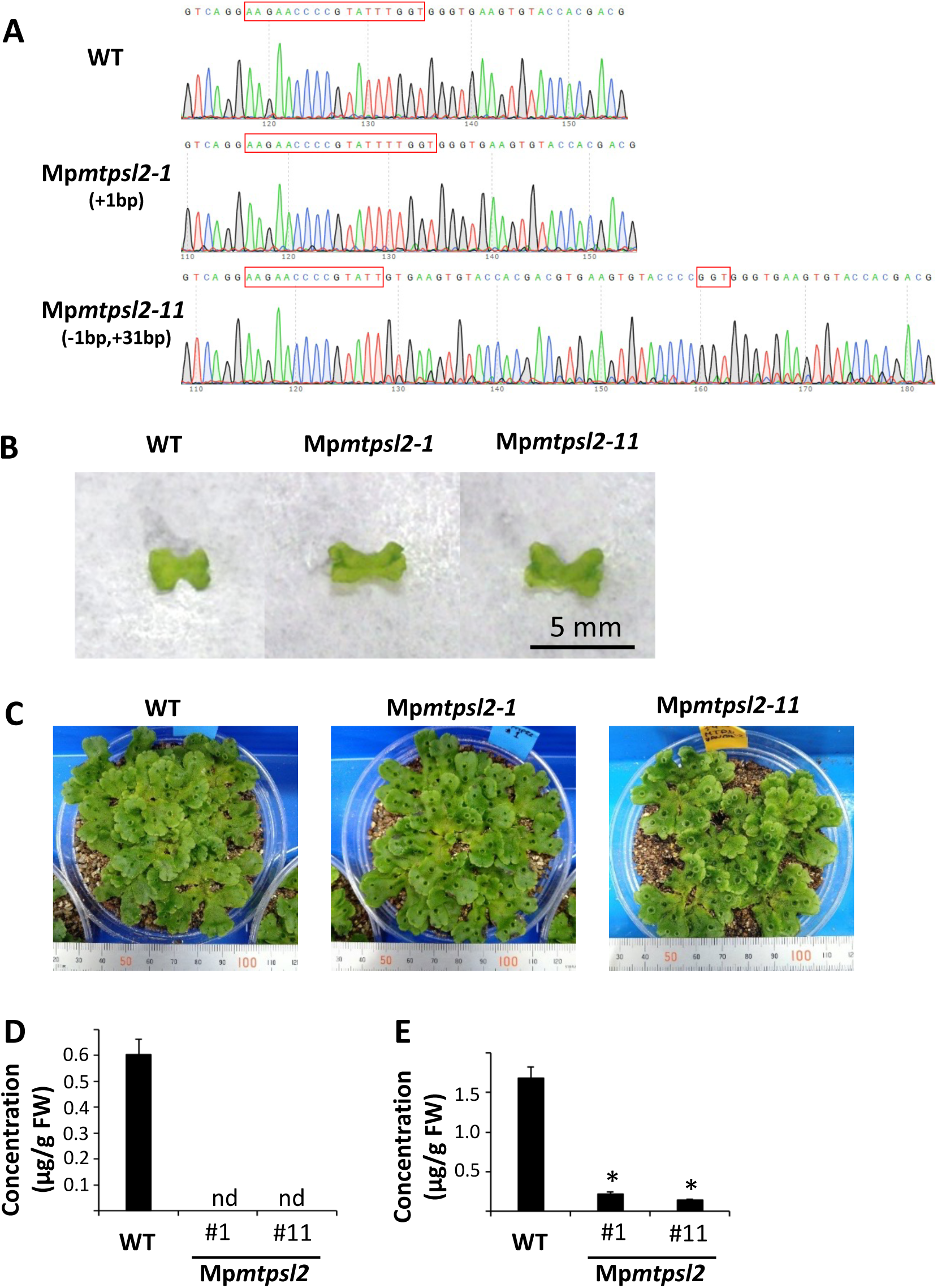
Production and analysis of loss-of-function Mp*mtpsl2* mutant plants. **A)** Confirmation of loss-of-function Mp*mtpsl2* mutations in the transgenic *M. polymorpha* lines Mp*mtpsl2-1* and Mp*mtpsl2-11*). WT stands for wild type plants. **B)** Phenotype of germinating gemmae of wild-type plants (WT) and two mutant lines Mp*mtpsl2-1* and Mp*mtpsl2-11*. **C)** Phenotype of mature wild-type plants (WT) and two mutant lines Mp*mtpsl2-1* and Mp*mtpsl2-11*. **D)** levels of α-Phellandrene in gemmae 6 days after germination from wild-type plants (WT) and two mutant lines Mp*mtpsl2-1* and Mp*mtpsl2-11*. **E)** levels of D-Limonene levels in gemmae 6 days after germination from wild-type plants (WT) and two mutant lines Mp*mtpsl2-1* and Mp*mtpsl2-11*. Significant differences were identified using one-way analysis of variance (*P* < 0.05, Dunnet tests). The asterisk indicates significant differences in the amount from that found with the wildtype. “nd” stands for not detected.

### Identification and functional determination of Mp*CPT5* as NPPS

With the demonstration that Mp*MTPSL2* is a major terpene synthase gene responsible for the biosynthesis of two monoterpenes α-phellandrene and D-limonene, the next question we asked is which gene is responsible for the production of its substrate. *In vitro* enzyme assays support that NPP is the substate of MpMTPSL2 (Fig. S1) (9). So far, NPPS has only been reported in a few species of flowering plants, where NPPSs are members of the CPT family (16). Here we hypothesized that in *M. polymorpha* NPP is also synthesized by a CPT enzyme. By searching for the *M. polymorpha* genome using CPTs from *Solanum lycopersicum* (tomato) as queries, eight CPT genes were identified. They were designated Mp*CPT1-8* (*SI Appendix*, Table S1). To narrow down the candidate *NPPS* gene, we performed co-expression analysis. Because Mp*CPT8* was annotated to encode a noncatalytic subunit, it was excluded. Using 504 publicly available RNA-seq libraries we addressed whether any Mp*CPT* genes were co-expressed with Mp*MTPSL2*. We observe Mp*CPT5* (Mp3g18510) is not only co-expressed (adjPValue = 4.02E-246) but also the top co-expression partner of Mp*MTPSL2* with a Pearson Correlation Coefficient (PCC) of 0.95 and a Mutual Rank (MR) of 1.73 (*SI Appendix*, Fig. S6).

To obtain functional evidence on the role of Mp*CPT5* as NPPS in *M. polymorpha*, we took a transgenic approach. Mp*CPT5* loss-of-function mutants were produced using CRISPR/Cas9. Two independent alleles, Mp*cpt5-1* and Mp*cpt5-7*, were identified and analyzed (Fig. 2A). The Mp*cpt5*-1 allele contains a 15 bp deletion in the first exon (*SI Appendix*, Fig. S7), whereas the Mp*cpt5-7* allele contains a 59 bp deletion in the first exon (*SI Appendix*, Fig. S8). As a result, the Mp*cpt5-1* allele and the Mp*cpt5-7* allele encode proteins of 340 amino acids and 58 amino acids, respectively, in contrast to the 345 amino acid protein encoded by the wild-type Mp*CPT5* (*SI Appendix*, Fig. S9). Under normal growing conditions, both germinating gemmae (Fig. 2B) and mature mutant plants (Fig. 2C) are morphologically similar to those of wild-type plants. We compared the concentrations of monoterpenes in geminated gemma (6 days after germination) of wild-type and mutant plants (Mp*cpt5-1* and Mp*cpt5-7*). In contrast to the wild type, neither α-phellandrene (Fig. 2D) nor D-limonene (Fig. 2E) was detected from the gemmae of either of the mutant lines.

**Figure 2.**
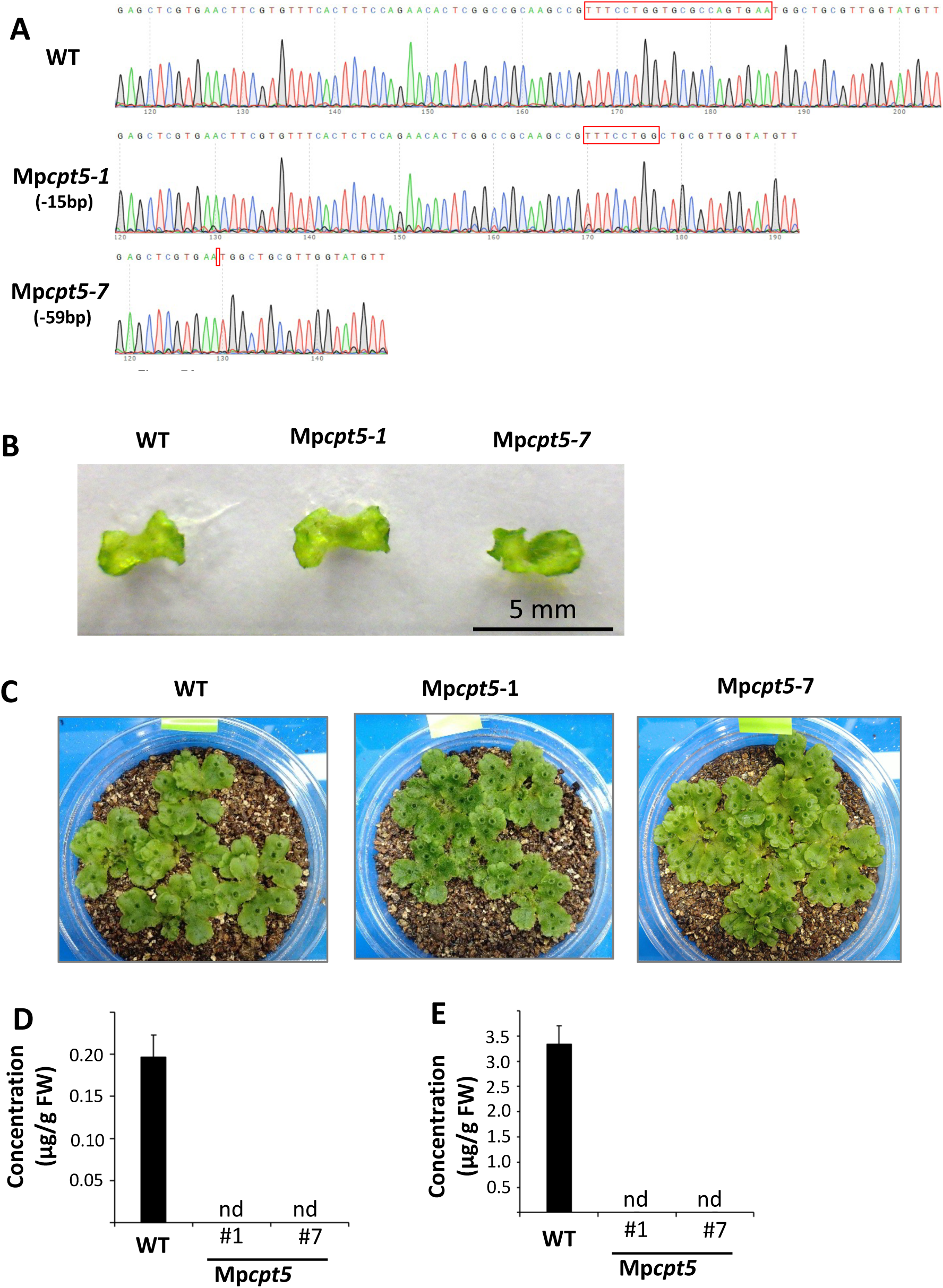
Production and analysis of loss-of-function Mp*cpt5* mutant plants. **A)** Confirmation of loss-of-function Mp*cpt5* mutations in the transgenic *M. polymorpha* lines Mp*cpt5-1* and Mp*cpt5-7*). WT stands for wild type plants. B) Phenotype of germinating gemmae of wild-type plants (WT) and two mutant lines Mp*cpt5-1* and Mp*cpt5-7*. **C)** Phenotype of mature wild-type plants (WT) and two mutant lines Mp*cpt5-1* and Mp*cpt5-7*. **D)** Levels of α-phellandrene in gemmae 6 days after germination from wild-type plants (WT) and two mutant lines Mp*cpt5-1* and Mp*cpt5-7*. **E)** Levels of D-limonene in gemmae 6 days after germination from wild-type plants (WT) and two mutant lines Mp*cpt5-1* and Mp*cpt5-7*. “nd” stands for not detected.

Mp*CPT5* was also analyzed for its catalytic activity using *in vitro* and *in vivo* approaches. For the in vitro approach, both full-length Mp*CPT5* and truncated Mp*CPT5* without the putative N-terminal transit peptide (*SI Appendix*, Fig. S10) were first cloned into protein expression vectors. Recombinant MpCPT5 proteins expressed in *E. coli* were subject to purification. However, they were found to be in inclusion bodies or to be unexpressed (*SI Appendix*, Fig. S11), preventing *in vitro* enzyme assays. For the *in vivo* approach, transient expression of Mp*CPT5* together with Mp*MTPSL2* in *Nicotiana benthamiana* was performed. Both α-phellandrene and D-limonene were detected (*SI Appendix*, Fig. S12). When Mp*CPT5* and Mp*MTPSL2* were transiently expressed alone, none of the two monoterpenes was detected (*SI Appendix*, Fig. S12).

### Both MpCPT5 and MpMTPSL2 are localized to the plastid

The conclusive demonstration that MpCPT5 and MpMTPSL2 catalyze sequential enzymatic steps of monoterpenes biosynthesis in *M. polymorpha* raises the question of whether monoterpene biosynthesis via NPP also occurs in the plastid, as do monoterpene biosynthesis pathways in seed plants. To address this, full-length Mp*CPT5* and Mp*MTPSL2* genes were each translationally fused to the yellow fluorescent protein (YFP) at their C-terminal end and then transiently expressed in protoplasts to determine subcellular locations. The result revealed that both MpCPT5 and MpMTPSL2 are localized in chloroplasts, but with different distribution patterns. While MpCPT5-YFP signals were detected in punctate structures that overlap with chloroplast autofluorescence, MpMTPSL2-YFP signals were found throughout chloroplasts (Fig. 3A).

**Figure 3.**
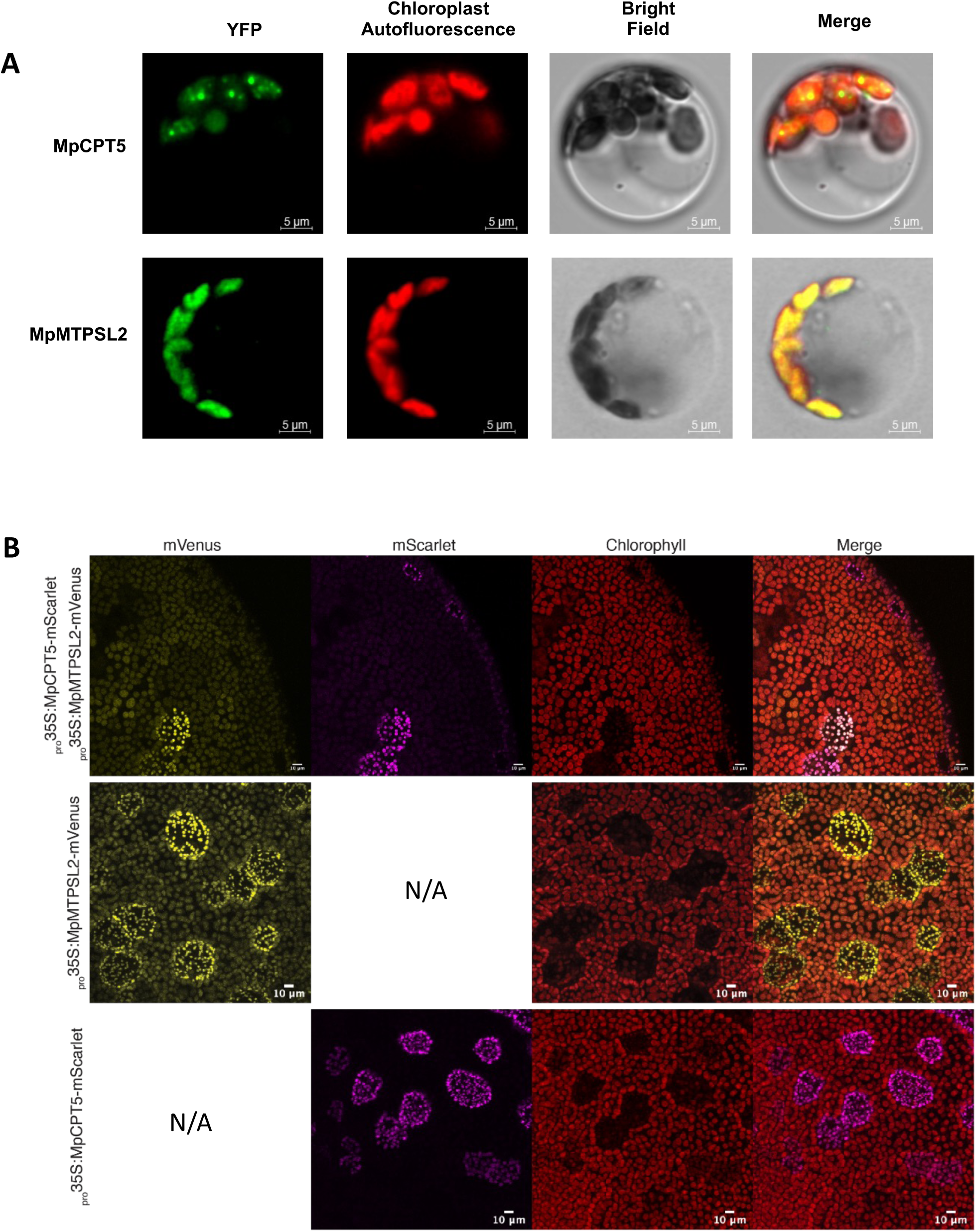
Subcellular localization of MpCPT5 and MpMTPSL2. **A)** Yellow fluorescence protein tagged full length MpCPT5 and MpMTPSL2 (MpCPT5-YFP and MpMTPSL2-YFP; green color) were expressed in Populus mesophyll protoplasts. The overlap of their fluorescent signals with chloroplast autofluorescence (red color) is indicated in yellow. Scale bars = 5 µm. **B)** subcellular localization of MpCPT5 and MpMTPSL2 in *M. polymorpha*. MpCPT5 and MpMTPSL2 are tagged with mScarlet (magenta) and mVenus (yellow), respectively. Transgenic *M. polymorpha* plants were produced individually with MpCPT5-mScarlet and MpMTPSL2-mVenus or via cotransformation of the two. Scale bars = 10 µm. “N/A” stands for not applicable.

To provide additional evidence on subcellular localizations of MpCPT5 and MpMTPSL2 in *M. polymorpha*, stable transgenic *M. polymorpha* plants expressing both Mp*MTPSL2* translationally fused to mVenus gene and Mp*CPT5* translationally fused to mScarlet gene under the control of a constitutive promoter (_pro_35S) were produced and analyzed. MpMTPSL2 and MpCPT5 were found to be co-localized in plastids of all cells with stronger signals in non-green plastids from oil body cells (Fig. 3B). As a control, transgenic *M. polymorpha* plants expressing either of the two expression cassettes were also produced. They showed the same result (Fig. 3B).

### Mp*CPT5* and Mp*MTPSL2* are specifically expressed in oil body cells

Oil bodies are specialized organelles only found in liverworts and contain specialized metabolites such as terpenes and bisbibenzyl compounds (22). It was shown previously that mutant plants lacking oil body cells are unable to accumulate sesquiterpenes and also D-limonene (23). Here we asked the question whether the biosynthesis of α-phellandrene and D-limonene catalyzed by MpCPT5 and MpMTPSL2 also occurs in oil body cells. Because high concentrations of α-phellandrene and D-limonene were observed in gemmae (Fig. S5), we examined the density and distribution of oil body cells in gemmae. MpSYP12B is an oil body-specific protein (25). Therefore, oil body cells can be visualized in transgenic *M. polymorpha* plants expressing a mCitrine (YFP)-MpSYP12B fusion gene under the control of MpSYP12B promoter (26). We examined gemmae of this transgenic plant and observed a relatively large number (approximately 30-60) of oil body cells around the gemma periphery (Fig. 4A), which is consistent with the high concentrations of α-phellandrene and D-limonene in gemmae.

**Figure 4.**
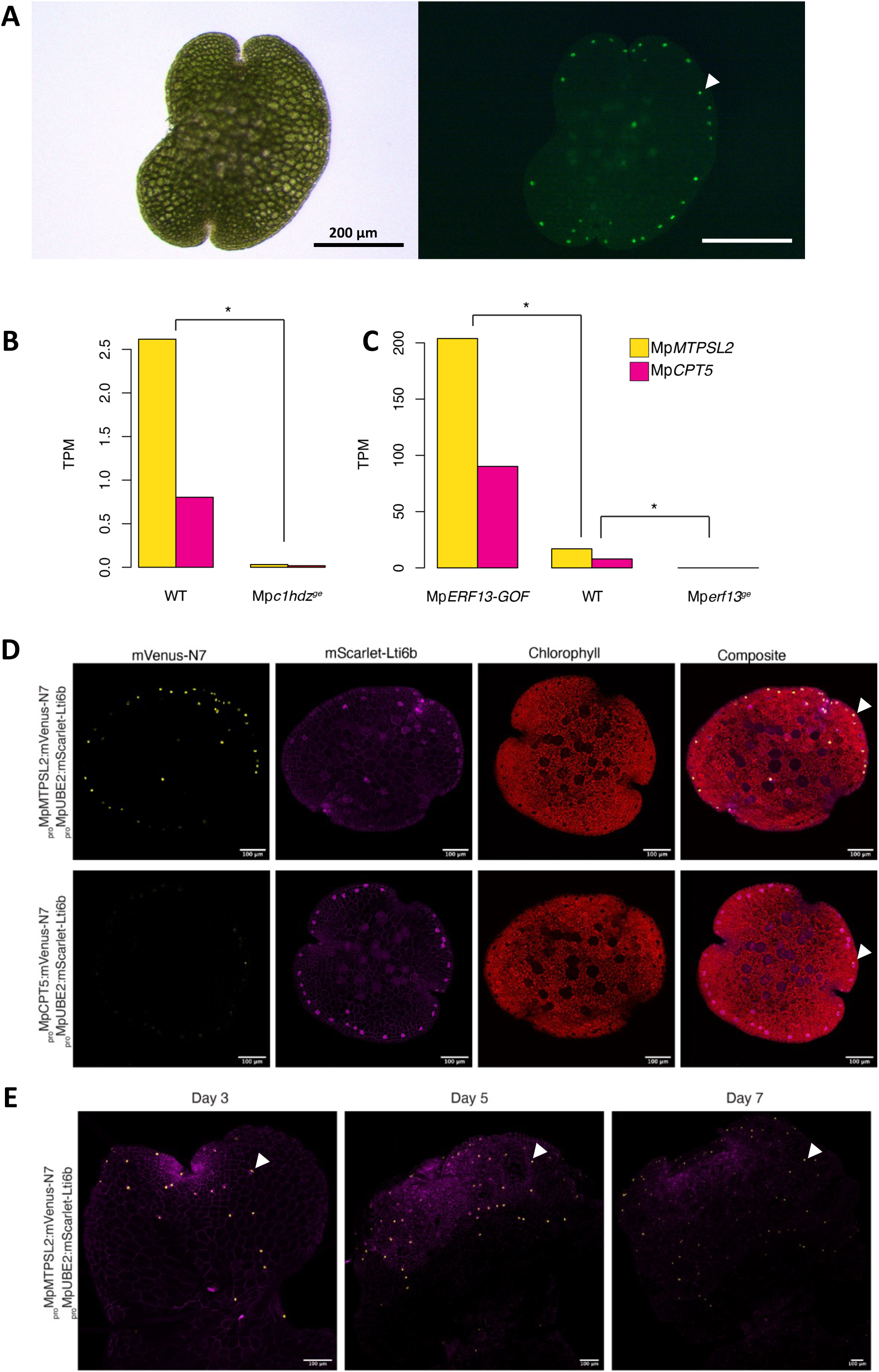
Cell-type specific expression of Mp*MTPSL2* and Mp*CPT5* to oil body cells. **A)** Relatively large number of (approximately 30-60) oil body cells on its periphery of the gemma. Oil body cells were identified by yellow fluorescence protein (mCitrine) which is fused to an oil body-specific protein. The fused gene was under the control of the promoter of Mp*SYP12B*. Scale bars = 200 µm. **B)** Expression of Mp*CPT5* and Mp*MTPSL2* in loss-of-function Mp*c1hdz* allele (Mp*c1hdz^ge^)* and wild-type plants (WT). **C)** Expression of Mp*CPT5* and Mp*MTPSL2* in loss-of-function Mp*erf13* allelle (Mp*erf13^ge^*), gain-of-function Mp*ERF13* (Mp*ERF13_GOF*) and wild-type plants (WT). TPM = transcripts per million. Asterisks indicate significant two-sided t-test (p-value < 0.01). **D**) Maximum projection of confocal images for the localization of promoter activity of Mp*MTPSL2* and Mp*CPT5* in oil body cells fused to nuclear localized mVenus-N7 (yellow). *_pro_*Mp*UBE2:mScarlet-Lti6b* (magenta) is a constitutive plasma membrane marker that also visualize the oil body membrane. Chlorophyll autofluorescence is visualized in red. Arrow indicates representative oil body cells. **E**) Time-course of _pro_Mp*MTPSL2* in 3-7 days old gemmalings. Arrow indicates representative oil body cells. Scale bars = 100 µm.

It has been established that oil body specification and differentiation requires Mp*ERF13* and Mp*C1HDZ*, both of which encode transcription factors (23, 26). In loss-of-function Mp*erf13* or Mp*c1hdz* alleles (23, 26), the expression of both Mp*CPT5* and Mp*MTPSL2* was found to be negligible compared to wild-type plants (Fig. 4B). In a gain-of-function Mp*ERF13* allele, expression of both Mp*CPT5* and Mp*MTPSL2* was significantly increased (Fig. 4C). Thus, expression of both Mp*CPT5* and Mp*MTPSL2* is correlated with the presence of oil body cells.

To test whether the genes are expressed in oil bodies, we constructed transcriptional fluorescent reporters for both Mp*CPT5* and Mp*MTPSL2* and examined expression during the first 7 days following gemma germination. As the genes are 1.5 kb apart from each other and transcribed in opposite directions, one promoter corresponds to the reverse complement of the other. As expected, the promoter activity in gemma of both Mp*MTPSL2* and Mp*CPT5* reporters is specific to oil body cells (Fig. 4D). Following gemma germination, _pro_MpMTPSL2 activity is lower in newly formed oil body cells and signal remains strong in oil bodies derived from the gemma (Fig. 4E), consistent with the accumulation of monoterpenes in gemma (Fig. S5). We conclude that the bidirectional promoter between both genes is sufficient to drive cell-type specific expression.

### Mp*CPT5* and Mp*MTPSL2* are located in a biosynthetic gene cluster

The genomic locus of Mp*MTPSL2* and Mp*CPT5* was identified as part of a putative biosynthetic gene cluster by PlantiSMASH (27). Plant biosynthetic gene clusters are characterized by H3K27me3 coverage and coordinated gene expression (28). We verified that this locus complies with both criteria and delimited the cluster to seven genes spanning approximately 35 kbp: Mp*MTPSL2*, Mp*CPT5*, four genes encoding hypothetical proteins, and Mp*CPT4* (Fig. 5A). Mp*CPT4* expression is at ten-fold lower levels compared to Mp*CPT5* and, as supported by genetics and co-expression studies, the functionally active enzymatic core of the cluster is limited to Mp*MTPSL2* and Mp*CPT5*, defining this locus as a potential biosynthetic gene cluster for monoterpene production.

**Figure 5.**
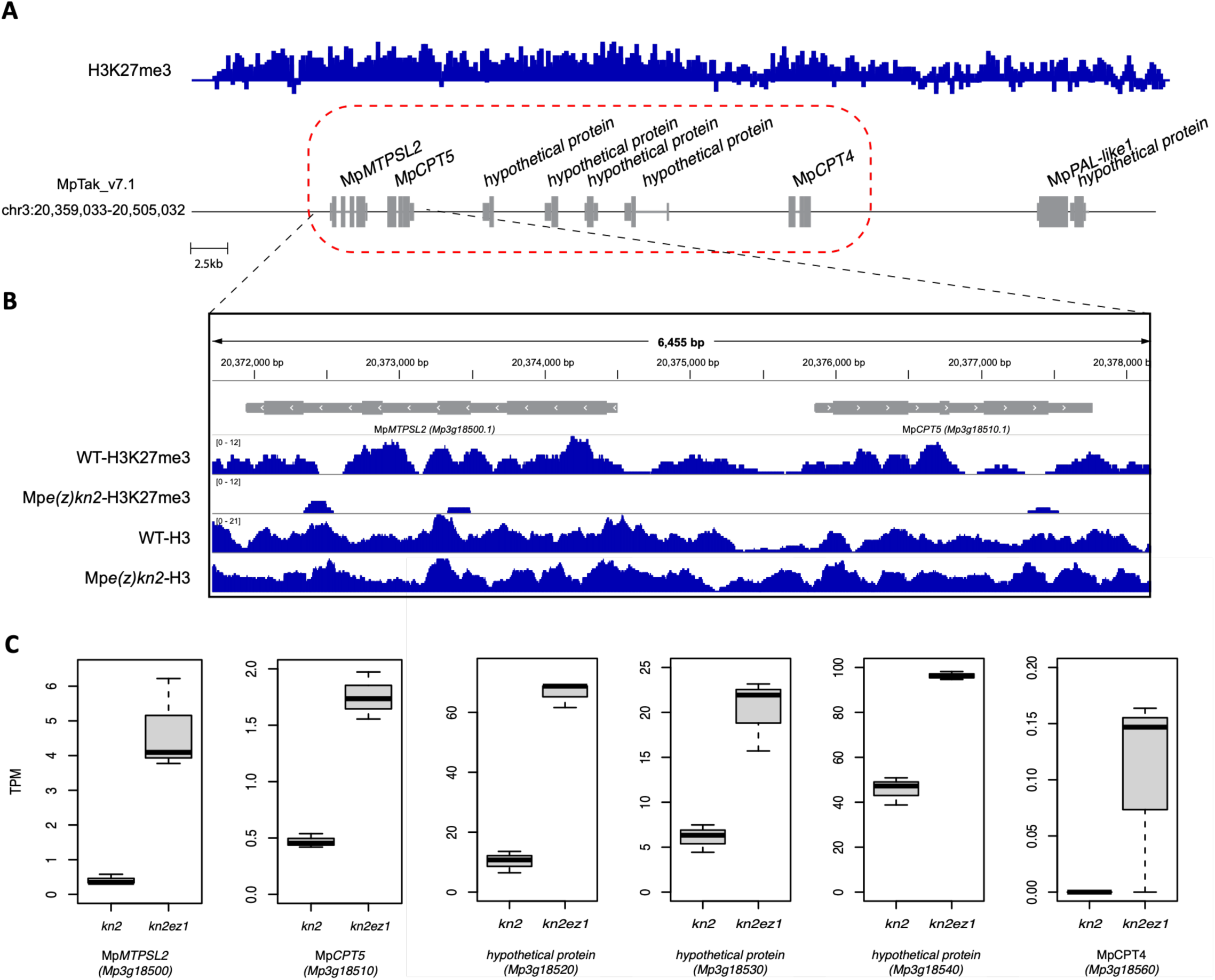
Mp*MTPSL2* and Mp*CPT5* operates as a biosynthetic gene cluster. **A)** Genomic overview of the chromosomal region containing the biosynthetic gene cluster on chromosome 3 (MpTak_v7.1). H3K27me3 CUT&RUN coverage is shown in blue from wild-type plants is shown above the gene annotation track (62). The dashed red box delineates the cluster according the epigenetic and co-expression landscape to (28), comprising Mp*MTPSL2* (Mp3g18500), Mp*CPT5* (Mp3g18510), four hypothetical protein-encoding genes (Mp3g18520, Mp3g18530, Mp3g18540, and Mp3g18550), and Mp*CPT4* (Mp3g18560). Scale bar, 2.5 kb. **B)** Integrative Genomics Viewer tracks showing H3K27me3 and total H3 ChIP-seq coverage over the MpMTPSL2–MpCPT5 gene pair in wild-type and Mpe(z)kn2 plants. Data are representative of two independent biological replicates. **C)** Expression levels (transcripts per million, TPM) of genes within the biosynthetic gene cluster in kn2 and kn2ez1 backgrounds (n = 3 biological replicates per genotype). All genes significantly upregulated in kn2ez1 relative to the kn2 control. Expression of Mp3g18550 was undetectable in both backgrounds and is therefore not shown. Boxplots show the interquartile range; whiskers extend to 1.5× the interquartile range; significance was determined by two-sided t-test.

PRC2-mediated H3K27me3 deposition maintains biosynthetic gene clusters in a repressed chromatin state in Arabidopsis, and loss of PRC2 function leads to cluster de-repression (28, 29). To determine whether the Mp*MTPSL2–*Mp*CPT5* cluster is similarly regulated, we examined H3K27me3 distribution and gene expression in the PRC2 loss-of-function mutant Mp*e(z)* Mp*knox2* (30). H3K27me3 enrichment was markedly reduced across the cluster locus in Mp*e(z)* Mp*knox2* plants relative to Mp*knox2* controls, and all six cluster genes showing detectable expression were significantly upregulated (Fig. 5B–C). These results demonstrate that the Mp*MTPSL2*–Mp*CPT5* locus operates as an epigenetically regulated biosynthetic gene cluster.

### Evolutionary relatedness of MpCPTs and CPTs from other plants

With the identification of Mp*CPT5* as encoding NPPS in *M. polymorpha*, one interesting is how this gene has evolved and how it is related to NPPS from angiosperms. To answer this question, phylogenetic analysis of CPT genes from 30 green plants covering all major linages of green plants (*SI Appendix*, Table S2) were performed. These include one species of green algae, eight species of liverworts including *M. polymorpha*, six species of mosses, one species of hornwort, three species of lycophytes, five species of ferns, one species of gymnosperm and five species of angiosperms including tomato. All seven CPT proteins from tomato have been functionally characterized (16). Sl*CPT1* is NPPS. SlCPT2 synthesizes nerylneryl diphosphate (C_20_), SlCPT6 produces Z,Z-FPP (C_15_). In contrast, SlCPT4, SlCPT5, and SlCPT7 produce longer-chain polyprenyl products (C_25_–C_55_). SlCPT3 showed no detectable *in vitro* activity but functionally complemented a yeast dolichol biosynthesis mutant (16). RNAi suppression of tomato SlCPT3 reduced dolichol content by ∼60%, and this catalytic subunit was found to require the distantly related, non-catalytic partner protein, SlCPTBP (31). CPTs form three distinct clades, designated CPT-a, b and c, of deep origin among the Viridiplantae (Fig. 6). CPT-a consist of non-catalytic subunit of CPTs, represented by SlCPTBP (Fig. 6). CPT-b contains known member of CPTs involved in dolicol biosynthesis (e.g., SlCPT3). CPT-c contains members of long-chain CPTs for the biosynthesis of polyprenl (SlCPT4, SlCPT5, and SlCPT7) and well as known short-chain CPTs (SlCPT1, SlCPT2 and SlCPT6). Mp*CPT5* is in the CPT-c clade. Within the clade of CPT-c, the phylogeny of CPT genes is congruent with species phylogeny. Mp*CPT5* is more closely related to other liverwort CPT genes than to SlNPPS genes of tomato (Fig. 6).

**Figure 6.**
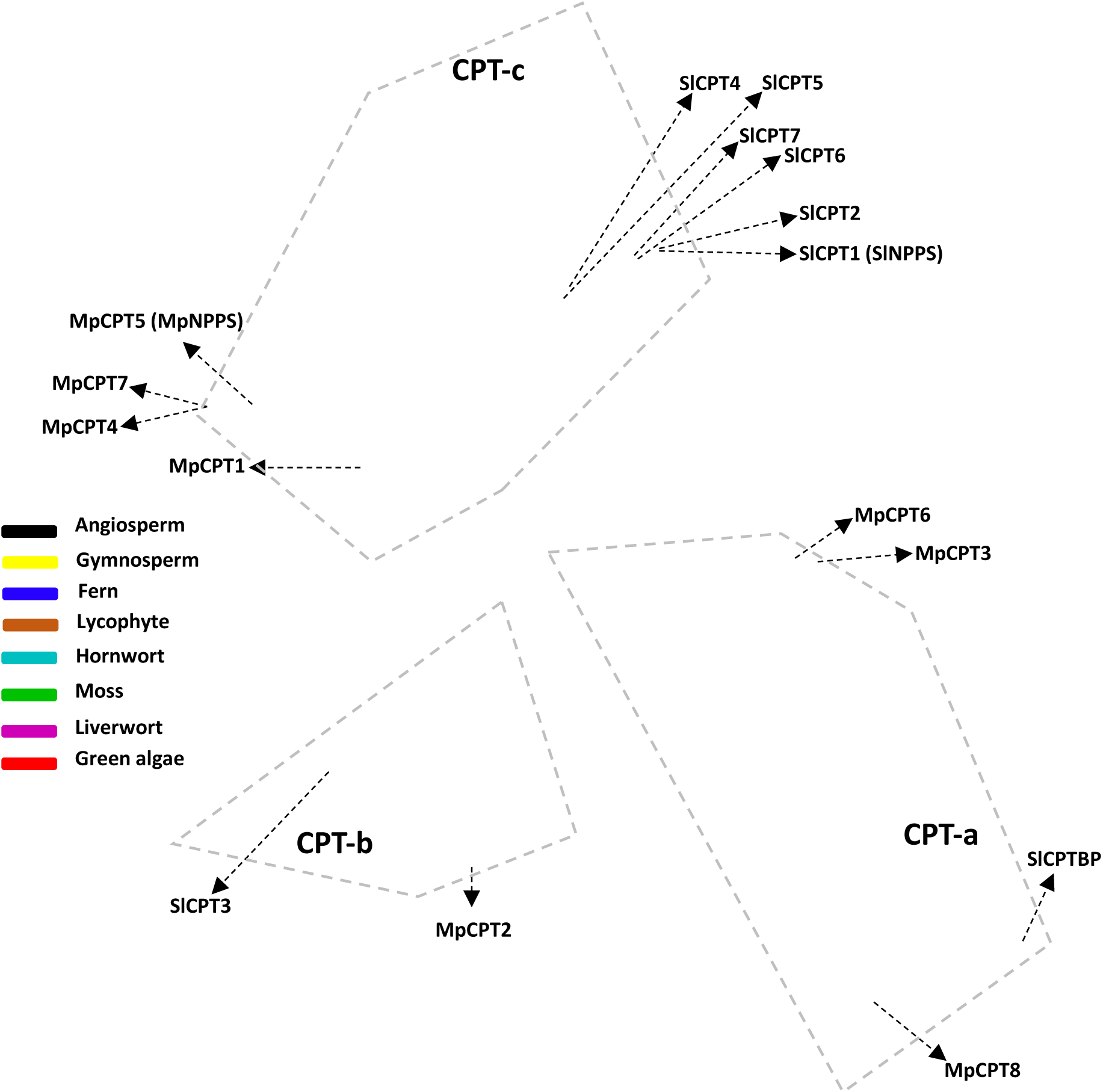
Phylogenetic tree of CPTs from *M. polymorpha* and other selected plants. The species included in this analysis are listed in *SI Appendix* Table S2. CPTs from various lineages of green plants are color-coded. CPTs from *M. polymorpha* are indicate as MpCTP1-8. CPTs from tomato are indicated as SlCPT1-7 and SlCPTBP.

## DISCUSSION

Geranyl diphosphate is the well-known substrate of monoterpenes in seed plants. In some flowering plants, NPP, the isomer of GPP, was demonstrated to serve as a complimentary substrate for monoterpene biosynthesis (5). In this work, we report that in the liverwort *M. polymorpha* NPP is the substrate for monoterpene biosynthesis. In particular, Mp*CPT5*-Mp*MTPSL2*, which encode NPPS and monoterpene synthase respectively, were determined to function as a module expressed in oil bodies to catalyze monoterpene biosynthesis. This biosynthetic module is defined based on three lines of evidence. Firstly, the product of MpMTP5, NPP, serves as the substrate for MpMTPSL2. As such, these two enzymes catalyze consecutive biochemical reactions. Secondly, Mp*CPT5* and Mp*MTPSL2* are physically linked in the genome sharing a bidirectional promoter. Thirdly, these two genes are co-regulated. These findings provide new insights into multiple topics pertaining to plant specialized metabolism, including the evolution of short-chain CPTs, general mechanism about monoterpene biosynthesis in land plants, the specific mechanisms underlying monoterpene biosynthesis in liverworts, and the mechanisms underlying coregulation of genes of a same biosynthetic pathway.

CPTs play an indispensable role in plant biology by catalyzing the biosynthesis of polyprenols and dolichols, the long-chain polyisoprenoid lipids that serve as membrane-anchored glycosyl carriers essential for N-linked protein glycosylation and cell wall polysaccharide biosynthesis (17). The presence of multiple *CPT* genes in plant genomes, whose products are targeted to the endoplasmic reticulum or chloroplast, underscores the importance of long-chain CPT activity in coordinating terpene metabolism across subcellular compartments. Loss of CPT function has been shown to result in severe developmental defects, highlighting the important role of these enzymes in plant growth and development (32). The presence of short-chain CPTs in the CPT gene family was a recent discovery. In tomato, Sl*CPT1*, encoding NPPS, is closed related to Sl*CPT2* and Sl*CPT6*, the other two short-chain CPTs (16). Because these three SlCPTs are clustered with three long-chain CPTs (Fig. 6), it can be inferred that these short-chain CPTs evolved from long-chain CPTs. Mp*CPT5* is the first short-chain CPT gene to be discovered from archegoniate plants. While the function of other CPTs from *M. ploymorpha* remains to be determined, it can be inferred that Mp*CPT5* evolved from long-chain CPT as well. Because the phylogeny of the CPTs in the CPT-c subfamily is largely congruent with the lineage phylogeny (Fig. 6), it can be concluded that the evolution of short-chain CPTs from long-chain CPTs in tomato and *M. polymorpha* occurred independently. It will be interesting to determine the catalytic functions of other CPT family members in *M. polymorpha* and CPTs from other liverworts. This information will help clarify whether short-chain CPTs are widespread in liverworts and whether they have deep evolutionary origins, since phylogenetic analysis alone (*SI Appendix*, Fig. S13) is insufficient to draw such conclusions. While it is unknown whether short-chain CPTs occur in other lineages of archegoniate plants, for example lycophytes and ferns, if they do, they would likely have evolved via independent evolution as well.

Our study demonstrates that biosynthesis of α-phellandrene and D-limonene is specific to oil body cells. This finding is consistent with the analysis on oil body cell mutants (23), co-expression analysis (26) and single-cell analysis performed in *Marchantia* (33). This also highlights that plastids contribute to the terpene repertoire of oil body cells. It was shown that oil body cell plastids are distinct from chloroplasts present in surrounding epidermal cells (24), suggesting that these plastids are differentiated into elaioplasts. Citrus peels accumulate terpenes and elaioplasts and chromoplasts of citrus peels are the sites of terpene biosynthesis, while terpenes specifically accumulate in elaioplasts (34). In many flowering plants, terpene biosynthesis occurs in specific secretory structures such as glandular trichomes (35). Despite oil body cells being unrelated to trichomes, they provide analogous compartments for the bioaccumulation of terpenes (36) and suggest another parallelism with the biosynthesis of monoterpenes in other flowering plant species such as *S. lycopersicum* (5). Recently, it has been reported that ABCG transporters localized to the oil body membrane are involved in the oil layer of oil body cells, specifically in the transport of specialized metabolites into oil bodies (37, 38). Monoterpenes like limonene also appear to be transported into the oil body lumen by oil body-specific ABCGs. This mechanism is similar to the transport of terpenoids into trichomes in flowering plants such as *Nicotiana* and *Mentha* (39).

In land plants, genes of the same biosynthetic pathways may have special arrangements on chromosomes to form biosynthetic gene clusters (BGCs), which are defined as groups of two or more non-homologous genes that are physically co-localized on the same chromosome and encode enzymes that catalyze consecutive steps in the same metabolic pathway. Mp*CPT5* and Mp*MTPSL2* meet this definition. Nevertheless, the *MpCPT5*-*MpMTPSL2* BGC is unique in that these two genes, of opposite orientation, are separated by 1,556 bp between their respective start codons (*SI Appendix*, Fig. S14). Our results demonstrate that the co-expression of these two genes (*SI Appendix*, Fig S6) as well as the cell-type specific expression is driven by a shared bidirectional promoter (Figs. 3 and 4). While such a mini-biosynthetic gene cluster with a bidirectional promoter involved in terpene biosynthesis has not been reported in plants, they have been discovered other organisms. For example, a terpene synthase-cytochrome P450 cluster with a bidirectional promoter has been discovered in the social amoeba *Dictyostelium discoideum* (40). The terpene synthase catalyzes the formation of a sesquiterpene discoidol, which is converted to a trisnorsesquiterpene named discodiene by the clustered cytochrome P450 (40). Despite the unique organization of Mp*CPT5*-Mp*MTPSL2* BGC among plants, the mechanism underlying its coregulation, specifically repression through H3K27me3 (Fig. 5), appears to be shared between this BGC and the typical BGCs preciously characterized (28). Biosynthetic gene clusters are often associated with cell-type or organ-specific biosynthetic gene pathways. For example, a gene cluster is responsible for the production of acyl-sugars in glandular trichomes of *S. lycopersicum* (41) and for triterpene synthesis in roots of *A. thaliana* and oat (42). In *S. lycopersicum*, the *TPS* and *NPPS* genes are located close to each other forming a functional cluster (43). This is now extended to the liverwort *M. polymorpha* because the Mp*CPT5*-Mp*MTPSL2* cluster exhibits specificity in oil body cells (Fig. 4). Overall, biosynthesis of monoterpenes in *M. polymorpha* is an extraordinary case of convergent evolution across land plant lineages, where co-regulation of enzymes, compartmentalization, and cell-type specificity are recurrent features.

The findings in this study are expected to inspire a number of new endeavors. It will be interesting to determine whether GPP is the primary substrate and NPP is a secondary substrate for monoterpene biosynthesis in archegoniate plants, as is the case in seed plants. An alternative hypothesis is that NPP is the primary substrate. Regardless of the scenario, it will be interesting to determine how the substrate, either GPP or NPP, is produced. It remains to be determined whether some archegoniate plants have evolved dedicated GPPS. For the production of NPP, it will also be interesting to determine whether NPPS is a conserved gene in some lineages. Regarding the unique biosynthetic gene cluster that we identified, it will be interesting to determine how widespread such mini-clusters with directional promoters are in plants, the mechanisms underlying their formation, and the mechanisms of gene coregulation, for instance, whether a single transcription factor can bind the bidirectional promoter to activate both genes.

## MATERIALS AND METHODS

### Plant material, growth conditions and chemicals

*M. polymorpha* L. *ruderalis* strains Tak-1 (male) (44) was used in this study. The gemmae were aseptically grown on half-strength Gamborg B5 medium (Fujifilm Wako Pure Chemicals, Osaka, Japan) solidified with 1% agar in a plastic plate (90 × 20 mm; Sansei Medical, Kyoto, Japan) at 22°C under continuous fluorescent light (36-61 μmol·m^−2^·s^−1^). Vermiculite was kept moist by regular watering. After supplying diluted Hyponex solution for the first time, no other nutrients were supplied. DMAPP, IPP, GPP and NPP were purchased from Echelon Biosciences (Salt Lake City, UT).

### Sequence retrieval and analysis

*CPT* genes were identified from the sequenced genomes of respective plants (*SI Appendix*, Table S2) using an HMMER search (45) with an e-value threshold of 1e-^5^. HMM profiles include PF19086 and PF03936 for MTPSL, PF01255 for CPT and PF00348 for IDS. After retrieving the genes, they were further verified using the NCBI Conserved Domain Search (https://www.ncbi.nlm.nih.gov/Structure/cdd/wrpsb.cgi). Multiple sequence alignment was performed using MAFFT (46) (L-INS-I) with 1000 iterations for improvement. The best-fit model of protein evolution was identified using ProtTest (47) under the Akaike Information Criterion. The maximum likelihood tree was constructed using RAxML with 1000 bootstrap replicates under the best substitution models (WAG+I+G for CPTs; JTT+G+F+I for MTPSLs). The tree was visualized using iTOL (https://itol.embl.de/).

### Transcriptomic and co-expression analysis

A TPM (transcripts per million) matrix from publicly available Marchantia libraries were obtained from MarpolBase Expression (https://mbex.marchantia.info/downloads) and the matrix was further reduced to select exclusively transcriptomic data (according to SRA Run and Library sources). Wild-type tissues were exclusively selected to track TPM values (above 1) and any libraries composed of mutant, pharmacological or environmental stress treatments were excluded from the analysis. The top five expressing libraries were selected using the slice_max (48) function. Plots were made using ggplot2 (49) in R Studio Version (2024.09.1+394). Raw reads were plotted using Integrative Genome Viewer (IGV 2.19.1) after mapping against MpTak_v7.1 (50) as described previously (51).

Co-expression analysis was made in R Studio using the cor(x,y, method = “pearson”) function to calculate PCC values and the cor.test(x,y, method = “pearson”)$p.value function to obtain P-Values from the correlation test. The Mp3g18500 TPMs were probed across the entire log2(X+1) transformed TPM matrix obtained from Marpolbase. Mutual rank values were calculated using reciprocal correlation tests and a formula previously described (52). GO-enrichment of Mp*MTPSL2* co-expressed genes as well as the overlap of co-expressed genes between Mp*MTPSL2* and Mp*CPT5* analysis was made using rank tables with the top 750 MR partners as previously described (52). For Mp*MTPSL2* and Mp*CPT5* expression in oil body mutants (23, 26), TPM values were obtained from MarpolBase Expression (52).

### Terpene synthase enzyme assays

*MpMTPSL2* was inserted into the pEXP5/CT-TOPO protein expression vector (Thermo Fisher Scientific, USA). Primers were summarized in *SI Appendix* Table S3. Using *E. coli* strain BL21 (DE3) pLysS (Life Technologies), recombinant proteins were produced by transformation of these constructs. For MTPSL enzymatic assay, protein production in *E. coli* and terpene synthase activity assays were conducted using NPP as a substrate, following previous established protocols for various isoprenyl diphosphate substrates (6, 21). Bacterial cultures expressing MpMTPSL2 cDNA were grown to an OD_600_ of 0.4, induced with 0.5 mM IPTG, and incubated at 16 °C for overnight. Cells were lysed using sonication in a chilled extraction buffer, and debris was removed via centrifugation. The supernatant was desalted into an assay buffer (10 mM Tris·HCl, pH 7.5, 1 mM DTT, 10% (vol/vol) glycerol). Enzyme activity was tested using bacterial extracts, NPP, 10 mM MgCl_2_, and 0.05 mM MnCl_2_ in sealed vials. Reaction products were collected using solid-phase microextraction fibers (SPME) and analyzed directly by Gas chromatogram-mass spectrometry (GC-MS).

### Expression and purification of recombinant MpCPT5

Full-length cDNA for MpCPT5 was cloned into two protein expression vectors pEASY-Blunt-E2 and pCzn1. A partial cDNA for truncated MpCPT5 (tMpCPT5) without the sequence coding for a putative N-terminal transit peptide with 53 amino acids (Fig. S10) was cloned into pET32a.

pEASY-Blunt E2-Mp*CPT5* and pET-32a-tMp*CPT5* were cotransformed with pG-KJE8 (Takara-Bio, Kyoto, Japan) for the expression of chaperone proteins into *E. coli* BL21(DE3). The strains BL21(DE3)/pG-KJE8/pEASY-Blunt E2-Mp*CPT5* and BL21(DE3)/pG-KJE8/pET-32a-tMp*CPT5* were cultivated at 37°C in 200 mL LB medium containing 100 mg/L ampicillin, 20 mg/L chloramphenicol, 0.5 g/L L-arabinose (for the expression of chaperone proteins), and 5 µg/L tetracycline (for the expression of chaperone proteins). The expression of MpCPT5 and tMpCPT5 was induced by adding 0.1 mM IPTG when OD_600_ reached 0.4, and the cells were cultivated overnight at 22°C. pCzn1-Mp*CPT5* was transformed into *E. coli* Top10 and the strain Top10/pCzn1-Mp*CPT5* was cultured in 200 mL LB medium containing 100 mg/L ampicillin at 37°C. When OD_600_ reached 0.4, 0.1 mM IPTG was added to induce expression, and the cells were cultivated overnight at 11°C. The cells were harvested by centrifugation and then suspended in 5 mL of 20 mM Tris-HCl buffer, pH 7.4, containing 500 mM NaCl and 20 mM imidazole per 1 g wet cells, followed by ultrasonic disruption. After centrifugation at 22,000×g at 4°C for 30 min, the supernatant and pellet fractions were analyzed by 12% SDS-PAGE.

### Transient expression in *N. benthamiana* leaves

Full-length cDNAs of *MpCPT5* and *MpMTPSL2* were cloned into the AgeI and XhoI restriction sites in the direct orientation of the pEAQ-HT binary vector (53). In the final construct, both *MpCPT5* and *MpMTPSL2* are under the control of the CaMV 35S promoter. *AtDXS2* was cloned into the XbaI and SacI sites in the direct orientation of pBI121, which is driven by the CaMV 35S promoter. *Agrobacterium tumefaciens* strain LAB4404 infiltration (agroinfiltration) was performed as described previously (54). Briefly, *A. tumefaciens* strain LBA4404 was grown at 28°C at 200 rpm for 24 h in Luria-Bertani medium containing kanamycin (50 mg L^−1^), rifampicin (50 mg L^−1^). Cells were collected by centrifugation for 15 min at 3,000g at 20°C and then resuspended in 10 mM MES buffer containing 10 mM MgCl_2_ and 100 μm acetosyringone (4′-hydroxy-3′,5′-dimethoxyacetophenone) to a final OD_600_ of 0.4, followed by incubation in the dark at 20°C for 3 h. For co-infiltration, equal numbers of cells from each of the cultures of strains harboring different binary plasmids were mixed together, collected by centrifugation, and resuspended as above to a final OD_600_ of 0.4 per strain. *Nicotiana benthamiana* plants were grown from seeds on soil in a greenhouse with a 16/8-h day/night photoperiod at 25°C. Leaves of 4-week-old *N. benthamiana* plants were infiltrated using a 2-mL syringe without a needle. The infiltrated leaves were collected 5 d after infiltration. The suppressor P19 from tomato bushy virus was co-expressed in all infiltrations to prevent post-transcriptional gene silencing and enhance transient expression in *N. benthamiana* (55). To analyze the compounds produced in the leaves, the harvested plant materials were flash frozen and ground into a fine power in liquid nitrogen. A 200 μL aliquot of 20% (v/v) NaCl was added to a 2 mL GC vial containing 0.2 g of plant tissue. The vial was placed on a 50°C hot plate, and volatiles were collected by SPME for 2 h. Analysis of the samples was performed with an Rxi-5Sil column on a Shimadzu GCMS-QP2010SE GC-MS system.

### GC-MS analyses

Gemmae collected from matured thalli, thalli grown from gemmae for 6, 8, 10 days on the agar plate, or the thalli grown for 2 weeks on vermiculite after transplanting from agar plate were used for volatile analysis. The gemmae or thalli (∼100 mg fresh weight) were collected in a screw-cap tube (2 mL) and snap-frozen in liquid nitrogen with eight stainless steel beads (3 mm i.d.). After addition of methyl *tert*-butyl ether (0.4 mL) containing 2.0 µg·mL^-1^ tetralin as the internal standard, the thalli were vigorously lysed with a beads cell disrupter (at 3,500 rpm for 1 min; Micro Smash MS-100R, Tomy, Tokyo, Japan). The mixture was centrifuged at 15,000 rpm for 3 min at 25°C, and the upper organic phase was directly subjected to GC-MS, essentially as described in Tanaka*, et al.* (56). In brief, a GC-MS system (QP-5050, Shimadzu, Kyoto, Japan) equipped with a DB-WAX column (30 m length × 0.25 mm i.d., 0.25 µm film thickness; Restek, Bellefonte, PA) was used. The column temperature was set as follows: 40°C for 5 min, increased by 5°C·min^-1^ to 200°C for 5 min for monoterpenes, or 80°C for 5 min, increased by 10°C·min^-1^ to 240°C for 5 min for sesquiterpenes. The carrier gas helium was delivered at a flow rate of 40 cm·s^-1^. The temperature of the injector and interface was 200°C. Splitless injection was used with 1 min sampling time. The mass spectrometer was operated in the electron impact mode with an ionization energy of 70 eV. D-limonene (Fujifilm Wako Pure Chemicals) and α-phellandrene (Tokyo Chemical Industry, Tokyo, Japan) were used to assign the respective peak and for quantification by using calibration curves. The calibration curve for each monoterpene with the internal standard (tetralin) was constructed to quantify its amount.

### Production of loss-of-function mutants of *M. polymorpha* for *MpMTPSL2* and *MpCPT5*

The gRNAs (*SI Appendix*, Table S3) were designed with Mp*MTPSL2* and Mp*CPT5* genes and cloned into the pMpGE_En03 binary vector carrying Cas9 to generate the CRISPR/Cas9 construct in pMpGE010 (57). The CRISPR/Cas9 vectors were transformed into *Agrobacterium tumefaciens* strain LBA4404 (for Mp*CPT5*) or GV2260 (for Mp*MTPSL2*). The CRISPR/Cas9 construct was further introduced into the thalli of *M. polymorpha* (Tak-1) according to the method previously reported (57). DNA regions near the target sequence of gRNA were amplified and directly sequenced.

### Subcellular localization of MpMTPSL2 and MpCPT5

Protoplast isolation and transfection were performed as previously described (58). The cDNAs of MpCPT5 and MpMTPSL2 were cloned into the pENTR/D-TOPO vector (Invitrogen) and then subcloned into the transient expression vector pUC-pGWB505 via LR reaction (Invitrogen) for C-terminal yellow fluorescent protein (YFP) fusion (58). For subcellular localization, 10 µL of MpCPT5-YFP and MpMTPSL2-YFP constructs were transfected into 100 µL of protoplasts, respectively. After 14 h incubation under weak light at room temperature, protoplasts were collected by centrifuge to subject to microscopy. Images were collected using a Zeiss LSM 710 confocal microscope, equipped with 514 and 561 nm laser lines for excitation of YFP and chloroplast autofluorescence, respectively. Images were processed using Zen software (Zeiss). Primers used are listed in *SI Appendix* Table S3.

For subcellular localization in stable transgenic plants of *M. polymorpha*, the CDS of MpMTPSL2 and MpCPT5 were synthesized by Genewiz. 35S promoter, mVenus or mScarlet, and the NOS terminator L0 parts were obtained from the OpenPlant toolkit (59). Parts were introduced into the pBy10 vector or pBy20, respectively (60), by one-step Type-IIS cloning using BsaI and T4 ligase. Plasmids were transformed into *A. tumefaciens* strain GV3101, which was then used to individually transform or co-transform *M. polymorpha*. Transgenic *M. polymorpha* plants were then selected for Hygromycin (pBy10), Chlorosulfuron (pBy20) or both.

### Generation of transcriptional reporters for MpMTPSL2 and MpCPT5

To obtain the mVenus fusion of the promoters, the intergenic region between MpMTPSL2 and MpCPT5 start codons was amplified with listed in Supplemental *SI Appendix* Table S3 and introduced into the binary vector pBy01, which contains a marker for Hygromycin selection *in planta* and a constitutive plasma membrane fluorescent marker (_pro_MpUBE2:mScarlet-Lti6b), and transformed into *M. polymorpha* sporelings as described in a recent study (61). Gemmalings were grown in contact plates for up to 7 days and imaged in a Leica SP5 confocal equipped with an Argon ion gas laser with emitted wavelengths of 458, 476, 488 and 514 nm and a 561 DPSS laser. Imaging was conducted using a 10× air objective (HC PL APO 675 10×/0.40 CS2). Excitation laser wavelengths and emission fluorescence bandwidth windows were as follows: for mVenus (514 nm, 527– 552 nm), for mScarlet (561 nm, 595– 620 nm) and for chlorophyll autofluorescence (687– 739 nm).

### Biosynthetic gene cluster identification and epigenetic analysis

Plant Specialized Metabolite Analysis 2.0 (PlantiSMASH) (27) pre-calculated database was used to define putative biosynthetic gene clusters in *Marchantia polymorpha*. Cluster boundaries were further delimited using co-expression data from MarpolBase Expression (MBEX) (52); and H3K27me3 enrichment, following the clique-based criteria described in (28). H3K27me3 CUT&RUN data from wild-type plants were obtained from SRA (SRX6721860-2) (62) RNA-seq data from kn2 and Mpe(z)kn2 plants were obtained from GEO (GSE221631) (30) and H3K27me3 ChIP-seq from GEO(GSE302232) (63). All sequencing data were mapped to the MpTak_v7.1 reference genome and visualized using the Integrative Genomics Viewer (IGV v2.19.1). Boxplots of gene expression within the cluster were generated using base R.

## Data, Materials, and Software Availability

The sequence data in this article can be obtained from the National Center for Biotechnology Information under the following GenBank accession number: PV233716 (MpCPT5). All other data are included in the article and/or *SI Appendix*.

## Acknowledgements

We thank Dr. Takehiko Kanazawa (National Institute for Basic Biology, Japan) for sharing transgenic *M. polymorpha* plants expressing mCitrine (YFP)-MpSYP12B fusion gene under the control of MpSYP12B promoter. We also thank Drs. Jianyu Fu and Minta Chaiprasongsuk for material assistance. This project was supported by the USDA Hatch fund and an Innovation Grant from the University of Tennessee, Institute of Agriculture (to F.C.) and Biotechnology and Biological Sciences Research Council BB/W014173/1 and BB/F011458/1 (to J.H). F.R. is a Leverhulme Early Career Fellow (ECF-2023-534) funded by the Leverhulme Trust and the Isaac Newton Trust (23.08(f)). E.F.-S. and J.L.B. were supported by The Australian Research Council Centre of Excellence for Plant Success in Nature and Agriculture (CE200100015). This material is also in part based upon work at the Centre for Bioenergy Innovation supported by the U.S. Department of Energy, Office of Science, Biological and Environmental Research under Contract Number ERKP886. K.M. was supported by JSPS KAKENHI Grant Numbers 23K26833, 23K27197.

## Author contributions

F.C., K.M., J.H., and J.L.B. planned and designed the research, G.W., T.Ka., F.R., E.F-S., M.X., X.C., T.Ko., K.I., M.N., and H.H. performed experiments, all authors contributed to data analysis, F.C., G.W., F.R., and K.M. wrote the manuscript. All authors revised the manuscript and approved the final version.

The authors declare no conflict of interest.

## Notice

This manuscript has been authored by UT-Battelle, LLC under Contract No. DE-AC05-00OR22725 with the U.S. Department of Energy. The United States Government retains and the publisher, by accepting the article for publication, acknowledges that the United States Government retains a non-exclusive, paid-up, irrevocable, worldwide license to publish or reproduce the published form of this manuscript, or allow others to do so, for United States Government purposes. The Department of Energy will provide public access to these results of federally sponsored research in accordance with the DOE Public Access Plan (http://energy.gov/downloads/doe-public-access-plan).

## Supplementary data

The following materials are available in the online version of this article.

**Figure S1.** Catalytic activity of MpMTPSL2.

**Figure S2.** Sequence of Mp*mtpsl2-1* allele.

**Figure S3.** Sequence of Mp*mtpsl2-11* allele.

**Figure S4.** Multiple sequence alignment of MpMTPSL2, Mpmtpsl2-1 and Mpmtpsl2-11.

**Figure S5.** Concentrations of monoterpenes in germinating of wild-type plants.

**Figure S6.** Co-expression analysis of Mp*MTPSL2* and Mp*IDS* genes.

**Figure S7.** Sequence of Mp*cpt5-1* allele.

**Figure S8.** Sequence of Mp*cpt5-7* allele.

**Figure S9.** Multiple sequence alignment of MpCPT5, Mpcpt5-1 and Mpcpt5-7.

**Figure S10.** Sequence of full-length and truncated MpCPT5.

**Figure S11.** Protein expression and purification of MpCPT5.

**Figure S12.** Metabolic analysis of *N. benthamiana* leaves after agroinfiltration.

**Figure S13.** Phylogenetic tree of CPTs.

**Figure S14.** Chromosomal organization of *MpMTPSL2* and *MpCPT5*.

**Table S1.** CPT genes in *M. polymorpha*.

**Table S2**. List of plants analyzed for the CPT family

**Table S3.** Primers used in this study.

