## Supplementary figures and tables for "Neryl diphosphate-derived monoterpene biosynthesis via a biosynthetic gene cluster in the liverwort *Marchantia polymorpha*"

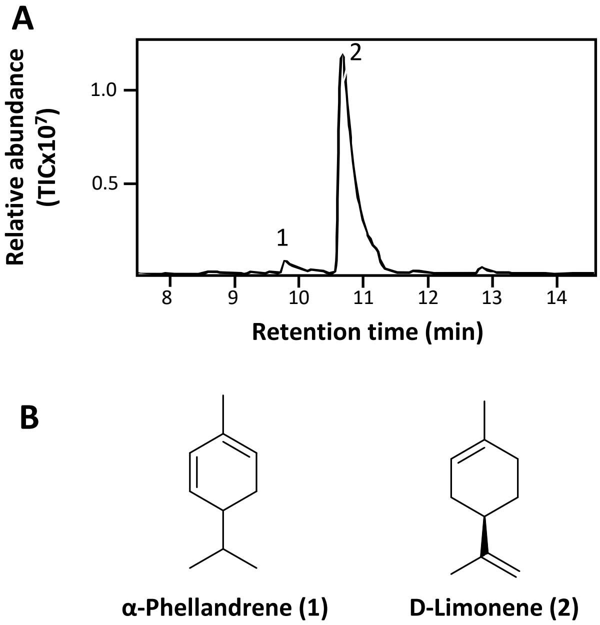

**Figure S1. catalytic activity of MpMTPLS2.** A) GC chromatogram of products of E. coli-expressed recombinant MpMTPSL2 using NPP as substrate. Peak 1 and peak 2 are identified as α-phellandrene (peak 1) and D-limonene (peak 2), respectively. B) Structure of α-Phellandrene (peak 1) and D-Limonene (peak 2).

**>MpMTPSL2 genomic seq**
**ATGGCGGCAAAAGCTCACAGGAGTTTCTCCAAGCTTATGAGTGTGAATTCGAGGTTCTTGTCCAGGAA**
**AAGCGAGGGCCAGAATGTCGCAGTGAAAGTGCCGATGCAGAATAATGTAGCATCCAAGGATCGAGTG**
**GTCGCCTTTTATGGCAGAGCAGCGTCACAGGCAAGGAGCCACCAAATTCAATCAGCAGCTTCATCAC**
**CTCAGCTGCCAGTTCCTCAGAAGGAGGAAAACCAAGTACCCGATTTCACTCATCTCTATGAGGACCGT**
**GGACTGAATCCTCTGATAGCCGAGTTCAATGAAAAGGCCCACGATTGGCTTGTCACGCACGATGTCC**
**ACACTCAGATGGGCCCGGAGGGATGGAAGGAATTCAAGTCAGGAAGAACCCCGTATTTTGGTGGGTG**
**AAGTGTACCACGACGTTGATCGGAAGGACTTTCTTTGGTGTGCCAAGTTTATAGCTTGGATTTCCTGTC**
**TTGATGATTTTTGCGACAAGCTTGAAATCTGGAGCGACATGAATAAAACCATCGGTCTCATCCTCGAA**
**ACGCAGAAGGTTATCTTGTGGTTCTGCTCTGAGGACCCTCGTCTCCTCCGAAACTTGGAGATGGTGTT**
**CGATTGTATCCCCACCGACCAACGTGACGATGTCTTTAGTGCATTTGCGGATTCCCTCGCCTACGCCC**
**GCACCAGCTCTCGCGCAG**GTGTTTCATTGATCTTAGTTCATGGGCTAGTCTCGCGTACACCTCCACGCA
GTCTTTGTGTTATTTAAATTTAAACGAGTTATCCAGGTCTGCATTCTGATAGCTATGGATCTCAAAACGAT
GACATCGTAAACTCTGACATAACTATCAATTCTGAAGCTGTCAATTAGTAGAAGTGGAGGTTAATATGTTA
TTCATTTCAGCTCTTCGACCTCTAATATTTTTTCAATGGTTGTTGCAG**GACTGATGAGTTTAGATTTGCAG**
**CCTTTAGCATCAGCTCTCAGAGAATTATTAATTGAGTACTGTGAACGCCTGCCATATGACTGCAATGTC**
**AGAATGGCCCAATATATACAGAATTATGCATTGACAATTGTTACTGAGACGAAGAGCAGATTATCCGC**
**AGAGTATCATACTTCAGTGGCTGAATTCACCGACTTTCGAACGCGATCCATTGGCGTGCAGCCCCTGA**
**TA**GTAAGCTTTCTTTTTCATTACGTCACGATTGCTTGCAGAATCGCGACCATGACATCCTAGACTCCATG
TCCTCTGCTTGCAAGAACGTGCTCACGTTATGAGCATCGATTGGCTGAATCGGGATCTGTCAGTCATGA
CGCGAAGGGACTTGTTGGTCTTAACTTCTGTCATATTTAAGGCTTGATTACTGCCATACGAGACTTTGGT
TCTTGACTTTAGTTTAGACTCTGGTTTAGCAGAATGAGCTGTAAACAATTGTCAACACGTTGCTGAAAGCT
GCTGAAATGCATTCAAAGAGTTCGACCACGTTGTGCATTTAGTATCATCTCAAGGTCCACAGTAGGTTCT
CAGCAGTCTTACGAGTTCAACCTTGTGCAG**GGTTTGACGGATCTCTTCATCGATTCTCGACGGTGGCCC**
**CAAATTCCGAAGGATGTGTTTGTCGATACAAACGTGCAAGCCCTCGTGGACGCTTCCTGCCGTGCCGT**
**AGCATGGGACAACGATATGTCCTCATTTTCAAAG**GTAAGAGGAAAATTTCCGCTGACAGTGCCATAGAC
ATAACAATATCACAAGAGTCCTATTATCAAGATTGTGTTTAGGCGAGTAGCTCCAGTCATTAGTGACGTG
TACCCCTTGGTTCCAATGTCTGTCGAGCTTTCCAGAAGTCCGTAGCAGCACTTTTATTTTGGTCCATTTC
CAAATTAACTGTGAAAGTCTTTATTTTGAAATTCTTTTCCAAAGTCTCCTGTGGTTTTTCTTTTAGCGCTAA
TTCAGTTTAATGTGAGCAAGGGATTGGTCACGGGCACCTAGGTCAGTATTGTCTCCAGAGGTAGGAAAC
AAAAAAGACTGCAATGGAATATTGAGAACTAGAACTTGATCTTTCTGGGGCACAGTCTTACAAAGGCTTT
ATGAGTGCCTTAG**GAGGTTCAAGAGCGAGGAGATCGTTTCAATGTGATCTTCATCATCATGGAAGAAT**
**GCGGATGTTCGCAAATCGAAGCTATGAATCGCGCGAAGGAAATGTTTCAAAACGAGATCAATGAGTT**
**CGAGACAAGAGTGGCAAACTTGAAGGTTGAACCTCAGTATCAGAATGCTGTGGCAGAGTACGTGAGA**
**ATATGCAGACTCATTATTCTAGGCATCCCACGATGGCATCGTAAGTCACAAAGATACCAAAAAGAGGA**
**CCACTACTTGAGATAA**

**Green: Exon**
**Black: Intron**
**Mp*mtpsl2*-1: T (+1 bp insertion)**

**Figure S2. Sequence of Mpmtpsl2-1 allele.**

**>MpMTPSL2 genomic seq**

**ATGGCGGCAAAAGCTCACAGGAGTTTCTCCAAGCTTATGAGTGTGAATTCGAGGTTCTTGTCCAGGAA**
**AAGCGAGGGCCAGAATGTCGCAGTGAAAGTGCCGATGCAGAATAATGTAGCATCCAAGGATCGAGTG**
**GTCGCCTTTTATGGCAGAGCAGCGTCACAGGCAAGGAGCCACCAAATTCAATCAGCAGCTTCATCAC**
**CTCAGCTGCCAGTTCCTCAGAAGGAGGAAAACCAAGTACCCGATTTCACTCATCTCTATGAGGACCGT**
**GGACTGAATCCTCTGATAGCCGAGTTCAATGAAAAGGCCCACGATTGGCTTGTCACGCACGATGTCC**
**ACACTCAGATGGGCCCGGAGGGATGGAAGGAATTCAAGTCAGGAAGAACCCCGTATT(t)GTGAAGTG**
**TACCACGACGTGAAGTGTACCCCGGTGGGTGAAGTGTACCACGACGTTGATCGGAAGGACTTTCTTTG**
**GTGTGCCAAGTTTATAGCTTGGATTTCCTGTCTTGATGATTTTTGCGACAAGCTTGAAATCTGGAGCGA**
**CATGAATAAAACCATCGGTCTCATCCTCGAAACGCAGAAGGTTATCTTGTGGTTCTGCTCTGAGGACC**
**CTCGTCTCCTCCGAAACTTGGAGATGGTGTTCGATTGTATCCCCACCGACCAACGTGACGATGTCTTT**
**AGTGCATTTGCGGATTCCCTCGCCTACGCCCGCACCAGCTCTCGCGCAG**GTGTTTCATTGATCTTAGTT
CATGGGCTAGTCTCGCGTACACCTCCACGCAGTCTTTGTGTTATTTAAATTTAAACGAGTTATCCAGGTC
TGCATTCTGATAGCTATGGATCTCAAAACGATGACATCGTAAACTCTGACATAACTATCAATTCTGAAGCT
GTCAATTAGTAGAAGTGGAGGTTAATATGTTATTCATTTCAGCTCTTCGACCTCTAATATTTTTTCAATGGT
TGTTGCAG**GACTGATGAGTTTAGATTTGCAGCCTTTAGCATCAGCTCTCAGAGAATTATTAATTGAGTA**
**CTGTGAACGCCTGCCATATGACTGCAATGTCAGAATGGCCCAATATATACAGAATTATGCATTGACAA**
**TTGTTACTGAGACGAAGAGCAGATTATCCGCAGAGTATCATACTTCAGTGGCTGAATTCACCGACTTT**
**CGAACGCGATCCATTGGCGTGCAGCCCCTGATA**GTAAGCTTTCTTTTTCATTACGTCACGATTGCTTGC
AGAATCGCGACCATGACATCCTAGACTCCATGTCCTCTGCTTGCAAGAACGTGCTCACGTTATGAGCAT
CGATTGGCTGAATCGGGATCTGTCAGTCATGACGCGAAGGGACTTGTTGGTCTTAACTTCTGTCATATTT
AAGGCTTGATTACTGCCATACGAGACTTTGGTTCTTGACTTTAGTTTAGACTCTGGTTTAGCAGAATGAG
CTGTAAACAATTGTCAACACGTTGCTGAAAGCTGCTGAAATGCATTCAAAGAGTTCGACCACGTTGTGCA
TTTAGTATCATCTCAAGGTCCACAGTAGGTTCTCAGCAGTCTTACGAGTTCAACCTTGTGCAG**GGTTTGA**
**CGGATCTCTTCATCGATTCTCGACGGTGGCCCCAAATTCCGAAGGATGTGTTTGTCGATACAAACGTG**
**CAAGCCCTCGTGGACGCTTCCTGCCGTGCCGTAGCATGGGACAACGATATGTCCTCATTTTCAAAG**GT
AAGAGGAAAATTTCCGCTGACAGTGCCATAGACATAACAATATCACAAGAGTCCTATTATCAAGATTGTG
TTTAGGCGAGTAGCTCCAGTCATTAGTGACGTGTACCCCTTGGTTCCAATGTCTGTCGAGCTTTCCAGAA
GTCCGTAGCAGCACTTTTATTTTGGTCCATTTCCAAATTAACTGTGAAAGTCTTTATTTTGAAATTCTTTTC
CAAAGTCTCCTGTGGTTTTTCTTTTAGCGCTAATTCAGTTTAATGTGAGCAAGGGATTGGTCACGGGCAC
CTAGGTCAGTATTGTCTCCAGAGGTAGGAAACAAAAAAGACTGCAATGGAATATTGAGAACTAGAACTTG
ATCTTTCTGGGGCACAGTCTTACAAAGGCTTTATGAGTGCCTTAG**GAGGTTCAAGAGCGAGGAGATCGT**
**TTCAATGTGATCTTCATCATCATGGAAGAATGCGGATGTTCGCAAATCGAAGCTATGAATCGCGCGAA**
**GGAAATGTTTCAAAACGAGATCAATGAGTTCGAGACAAGAGTGGCAAACTTGAAGGTTGAACCTCAG**
**TATCAGAATGCTGTGGCAGAGTACGTGAGAATATGCAGACTCATTATTCTAGGCATCCCACGATGGCA**
**TCGTAAGTCACAAAGATACCAAAAAGAGGACCACTACTTGAGATAA**

**Green: Exon**

**Black: Intron**

**Mp*mtpsl2*-11: t (-1 bp deletion),**
**GTGAAGTGTACCACGACGTGAAGTGTACCCC (+31 bp insertion)**

**Figure S3. Sequence of Mpmtpsl2-11 allele.**

```
MpMTPSL2      MAAKAHRSFSKLMSVNSRFLSRKSEGQNVAVKVPMQNNVASKDRVVAFYGRAASQARSHQIQSAASSPQLPVPQKEENQVPDFTHLYEDRGLNPLIAEFN   100
Mpmtpsl2-1    MAAKAHRSFSKLMSVNSRFLSRKSEGQNVAVKVPMQNNVASKDRVVAFYGRAASQARSHQIQSAASSPQLPVPQKEENQVPDFTHLYEDRGLNPLIAEFN   100
Mpmtpsl2-11   MAAKAHRSFSKLMSVNSRFLSRKSEGQNVAVKVPMQNNVASKDRVVAFYGRAASQARSHQIQSAASSPQLPVPQKEENQVPDFTHLYEDRGLNPLIAEFN   100
Consensus     maakahrsfsklmsvnsrflsrksegqnvavkvpmqnnvaskdrvvafygraasqarshqiqsaasspqlpvpqkeenqvpdfthlyedrglnpliaefn

MpMTPSL2      EKAHDWLVTHDVHTQMGPEGWKEFKSGRTPY..........LVGEVYHDVDRKDFLWCAKFIAWISCLDDFCDKLEIWSDMNKTIGLILETQKVILWFCS   190
Mpmtpsl2-1    EKAHDWLVTHDVHTQMGPEGWKEFKSGRTPY..........FGG........................................................   134
Mpmtpsl2-11   EKAHDWLVTHDVHTQMGPEGWKEFKSGRTPYCEVYHDVKCTPVGEVYHDVDRKDFLWCAKFIAWISCLDDFCDKLEIWSDMNKTIGLILETQKVILWFCS   200
Consensus     ekahdwlvthdvhtqmgpegwkefksgrtpy            g

MpMTPSL2      EDPRLLRNLEMVFDCIPTDQRDDVFSAFADSLAYARTSSRAGLMSLDLQPLASALRELLIEYCERLPYDCNVRMAQYIQNYALTIVTETKSRLSAEYHTS   290
Mpmtpsl2-1    ....................................................................................................   134
Mpmtpsl2-11   EDPRLLRNLEMVFDCIPTDQRDDVFSAFADSLAYARTSSRAGLMSLDLQPLASALRELLIEYCERLPYDCNVRMAQYIQNYALTIVTETKSRLSAEYHTS   300
Consensus

MpMTPSL2      VAEFTDFRTRSIGVQPLIGLTDLFIDSRRWPQIPKDVFVDTNVQALVDASCRAVAWDNDMSSFSKEVQERGDRFNVIFIIMEECGCSQIEAMNRAKEMFQ   390
Mpmtpsl2-1    ....................................................................................................   134
Mpmtpsl2-11   VAEFTDFRTRSIGVQPLIGLTDLFIDSRRWPQIPKDVFVDTNVQALVDASCRAVAWDNDMSSFSKEVQERGDRFNVIFIIMEECGCSQIEAMNRAKEMFQ   400
Consensus

MpMTPSL2      NEINEFETRVANLKVEPQYQNAVAEYVRICRLIILGIPRWHRKSQRYQKEDHYL                                                 444
Mpmtpsl2-1    ......................................................                                                 134
Mpmtpsl2-11   NEINEFETRVANLKVEPQYQNAVAEYVRICRLIILGIPRWHRKSQRYQKEDHYL                                                 454
Consensus
```

**Figure S4. Multiple sequence alignment of MpMTPSL2, Mpmtpsl2-1 and Mpmtpsl2-11.**

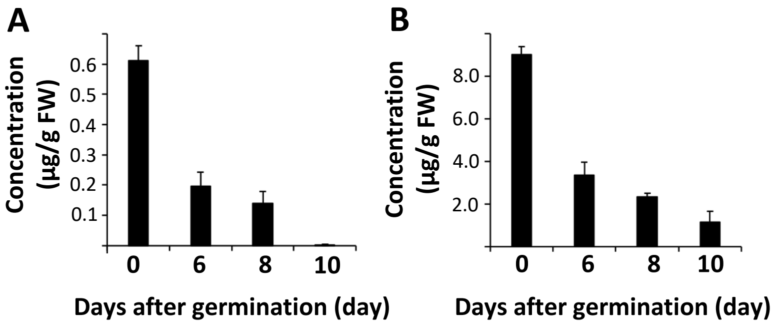

**Figure S5. Concentrations of monoterpenes in germinating of wild-type plants.** A) α-phellandrene levels in gemmae over time after germination. B) D-limonene levels in gemmae over time after germination.

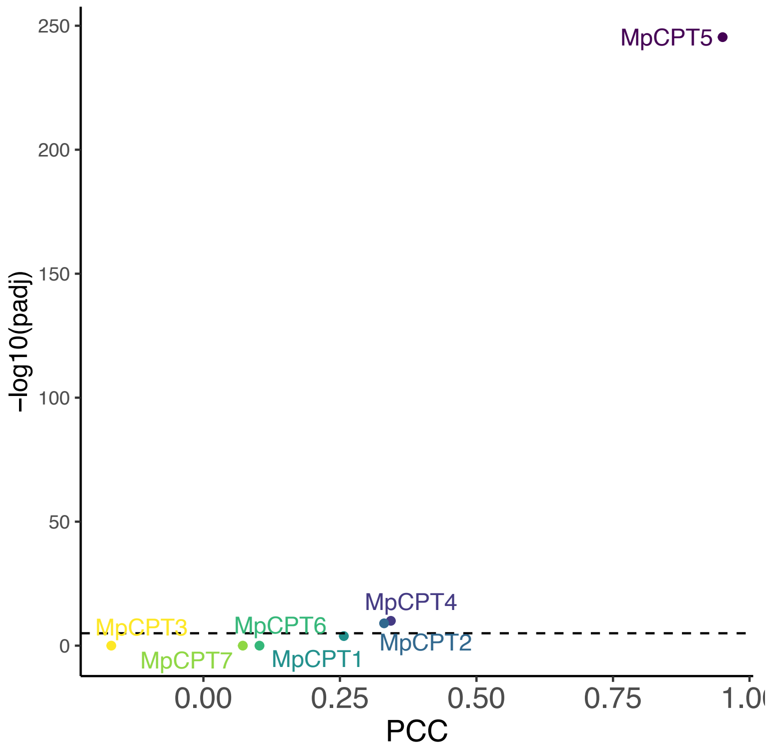

**Figure S6** Co-expression analysis of seven *MpCPT* genes against *MpMTPSL2*, x-axis shows PCC values and y-axis shows –log10 transformed P Values of Pearson correlation.

**>MpCPT5 genomic seq**

**ATGGAGTCTGTGGTGGCAAGATGTGTGAATCTGAAATCCGTACAATTGTCCAC**
**GATTTACAAGACGCATTGTGAAGCTGCAGCACCGATGGAGCTAAGCATGAGTA**
**GATCAACGATACCTGGCGCGAGCATCAGCAGTCACCGAGCTCGTGAACTTCG**
**TGTTTCACTCTCCAGAACACTCGGCCGCAAGCCGTTTCCTGG(TGCGCCAGTG**
**AATGG)CTGCGTTGGTATGTTGCATGAGACAGCCGGGACCGCATCAGCTGACA**
**GTTGTGCAGATCTGAGCCAGACGAAGAGGAGGATCGAACTTCCCCCTGAACT**
**GAATCCTGAGGTAATGCCCAAGCATGTTGCTCTTATCTTAGATGGCAACAATCG**
**TTACTCGAAGGCGAGAGGCGTGAGGGGAAGTGAAGGCTTCGAAGTTGGGCT**
**CAAGAAGAGTCTCAAAGACGCTGTGAGAGTCAGCGTTGATTGGGGAATCGAG**
**ATACTCACACTCTTTTACTTTTCTTACGACAATTGGAATCGAGAAAAG**GTTATTT
CTAAAACCCATTTCCGGTTATATTGCAGTTTAAAGAGCGTAACATGTTATGTCCGG
ACCCGTTTAGAGTTCTTCATGCTTAGGTGTGCCCAGAATATCGTCAAGGTGTTGT
AAGTTAGGTTCCAGTTTGATGACACTGCATGACCAGCGGTAGATGTGGACTTGA
TGGTTCTGAAGCGGAAGGTGTGATGGAGGTTGGATACAG**GTTGAAGTTGAAAC**
**AATATTTGGGCTTTTCGAGATATATCTGATCAAGTACAGAGAAGCCGTAATGAG**
GTCAGAAAACCTGGCTTTTGTTTCGTTATCTGTTTCAAATTCTCTAAAAGTACTGT
CCATATTCCTCCCTAATACTAGAAAAAATTTCATGATGGTTGAGTTAAAACAATCTA
TATTTTTGTATCTGTCAGTTACTATCTCTTGCGTAAATATCACAGCATTTGGCAATG
CCCTGCGGCCTGGCCTTGATGATCTATAACGAAAGTTTATACGCTGTTGATTGTA
CAG**GTATAACATTCAGTTTTCGGTCATGGGCACCCCAGAGAGATTTCCTGAGA**
**CTTTGCAAAAACTGGTGGCTAAGGTGACGGAAGAGTCTCGCAACAACAAGGG**
**ACTGAAGGTTGTTGTGGCAGCCGGGTACGGTGGAAGACAGGACATTCTTCAA**
**GCTACGCAAAGAATATCTCAACTGGTGGCTAACACAGAACTCTCCATCGATGA**
**GATAACACTGGAGGTGTTCGAGTCACACCTCATGACAAAAAACGTGAATCCAT**
**CCAGTGTTGATCTTATGATTCGAACAAGCGGAGAGCAGAGAATTAGCAACTTC**
**CTTCTATGGCAGACGGCGTGGACCGAGTTTGTATTTTTGAAGGAAAACTGGCC**
**CGAGTTTAGACGTGATTCGATGAAGAATGCGCTAATCAGTTACCAGTCTAGGG**
**ACCGAAGGCTTGGTCTAAGACAGACCAAGTTCTGA**

**Green: Exon**
**Black: Intron**
**Mp*cpt5*-1: TGCGCCAGTGAATGG (-15 bp deletion)**

**Figure S7. Sequence of Mpcpt5-1 allele.**

**>MpCPT5 genomic seq**

**ATGGAGTCTGTGGTGGCAAGATGTGTGAATCTGAAATCCGTACAATTGTCCAC**
**GATTTACAAGACGCATTGTGAAGCTGCAGCACCGATGGAGCTAAGCATGAGTA**
**GATCAACGATACCTGGCGCGAGCATCAGCAGTCACCGAGCTCGTGAA(CTTCG**
**TGTTTCACTCTCCAGAACACTCGGCCGCAAGCCGTTTCCTGGTGCGCCAGTGA**
**A)TGGCTGCGTTGGTATGTTGCATGAGACAGCCGGGACCGCATCAGCTGACAG**
**TTGTGCAGATCTGAGCCAGACGAAGAGGAGGATCGAACTTCCCCCTGAACTG**
**AATCCTGAGGTAATGCCCAAGCATGTTGCTCTTATCTTAGATGGCAACAATCGT**
**TACTCGAAGGCGAGAGGCGTGAGGGGAAGTGAAGGCTTCGAAGTTGGGCTC**
**AAGAAGAGTCTCAAAGACGCTGTGAGAGTCAGCGTTGATTGGGGAATCGAGA**
**TACTCACACTCTTTTACTTTTCTTACGACAATTGGAATCGAGAAAAG**GTTATTTC
TAAAACCCATTTCCGGTTATATTGCAGTTTAAAGAGCGTAACATGTTATGTCCGGA
CCCGTTTAGAGTTCTTCATGCTTAGGTGTGCCCAGAATATCGTCAAGGTGTTGTA
AGTTAGGTTCCAGTTTGATGACACTGCATGACCAGCGGTAGATGTGGACTTGAT
GGTTCTGAAGCGGAAGGTGTGATGGAGGTTGGATACAG**GTTGAAGTTGAAACA**
**ATATTTGGGCTTTTCGAGATATATCTGATCAAGTACAGAGAAGCCGTAATGAG**G
TCAGAAAACCTGGCTTTTGTTTCGTTATCTGTTTCAAATTCTCTAAAAGTACTGTC
CATATTCCTCCCTAATACTAGAAAAAATTTCATGATGGTTGAGTTAAAACAATCTAT
ATTTTTGTATCTGTCAGTTACTATCTCTTGCGTAAATATCACAGCATTTGGCAATGC
CCTGCGGCCTGGCCTTGATGATCTATAACGAAAGTTTATACGCTGTTGATTGTAC
AG**GTATAACATTCAGTTTTCGGTCATGGGCACCCCAGAGAGATTTCCTGAGACT**
**TTGCAAAAACTGGTGGCTAAGGTGACGGAAGAGTCTCGCAACAACAAGGGAC**
**TGAAGGTTGTTGTGGCAGCCGGGTACGGTGGAAGACAGGACATTCTTCAAGC**
**TACGCAAAGAATATCTCAACTGGTGGCTAACACAGAACTCTCCATCGATGAGA**
**TAACACTGGAGGTGTTCGAGTCACACCTCATGACAAAAAACGTGAATCCATCC**
**AGTGTTGATCTTATGATTCGAACAAGCGGAGAGCAGAGAATTAGCAACTTCCT**
**TCTATGGCAGACGGCGTGGACCGAGTTTGTATTTTTGAAGGAAAACTGGCCCG**
**AGTTTAGACGTGATTCGATGAAGAATGCGCTAATCAGTTACCAGTCTAGGGAC**
**CGAAGGCTTGGTCTAAGACAGACCAAGTTCTGA**

**Green: Exon**
**Black: Intron**
**Mp*cpt5-7*:**
**CTTCGTGTTTCACTCTCCAGAACACTCGGCCGCAAGCCGTTTCC**
**TGGTGCGCCAGTGAA (-59 bp deletion)**

**Figure S8. Sequence of Mpcpt5-7 allele.**

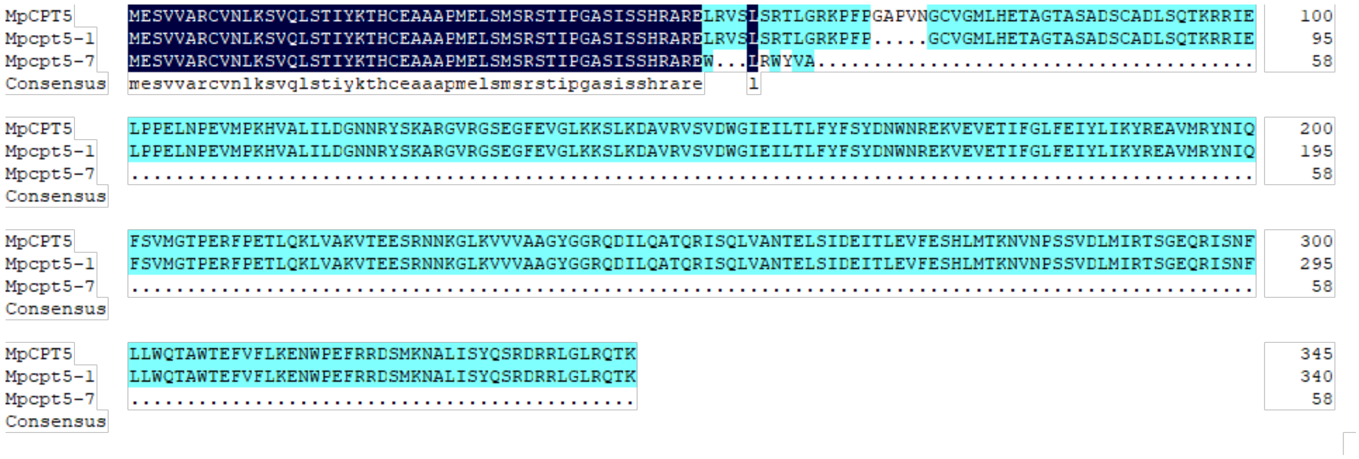

**Figure S9. Multiple sequence alignment of MpCPT5, Mpcpt5-1 and Mpcpt5-7.**

>MpCPT5

MESVVARCVNLKSVQLSTIYKTHCEAAAPMELSMSRSTIPGASIS
SHRARELRVSLSRTLGRKPFPGAPVNGCVGMLHETAGTASADSC
ADLSQTKRRIELPPELNPEVMPKHVALILDGNNRYSKARGVRGSE
GFEVGLKKSLKDAVRVSVDWGIEILTLFYFSYDNWNREKVEVETIF
GLFEIYLIKYREAVMRYNIQFSVMGTPERFPETLQKLVAKVTEESR
NNKGLKVVVAAGYGGRQDILQATQRISQLVANTELSIDEITLEVFE
SHLMTKNVNPSSVDLMIRTSGEQRISNFLLWQTAWTEFVFLKEN
WPEFRRDSMKNALISYQSRDRRLGLRQTKF*

**Figure S10. Sequence of full-length and truncated MpCPT5.** The first 53 amino acids in green indicate the putative transit peptide removed in the truncated version of MpCPT5.

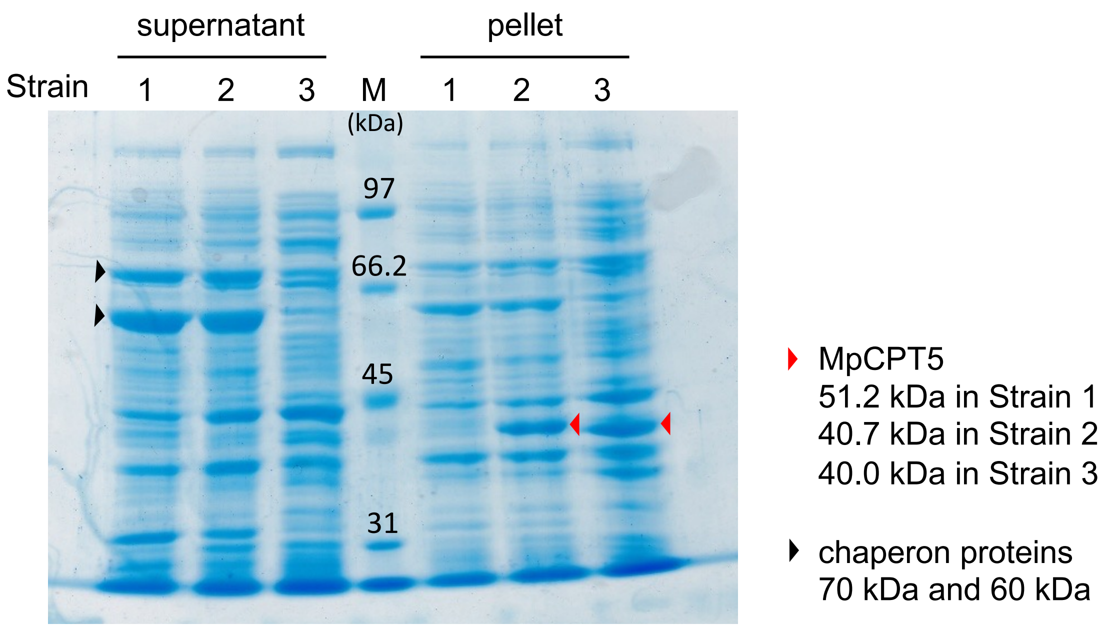

**Figure S11. SDS-PAGE analysis of the recombinant MpCPT5 expressed in *E. coli*.** Strain 1: BL21(DE3)/pG-KJE8/pET-32a-tMpCPT5 Strain 2: BL21(DE3)/pG-KJE8/pEASY-Blunt E2-MpCPT5. Strain 3: Top10/pCzn1-MpCPT5. M: marker proteins. Black and red arrowheads indicate the putative protein bands of chaperon proteins and MpCPT5s, respectively.

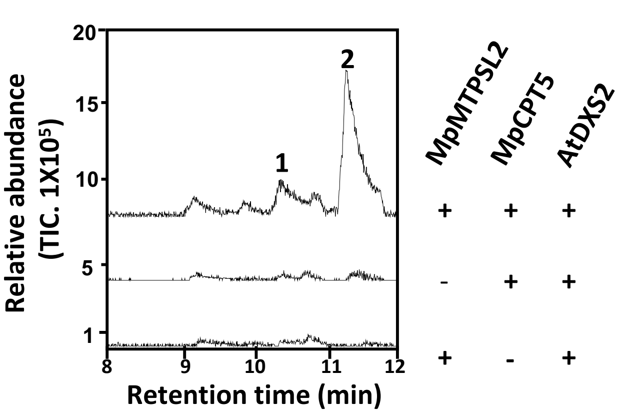

**Figure S12. Metabolic analysis of *N. benthamiana* leaves after agroinfiltration. *N. benthamiana* leaves transiently expressing MpCPT5 and MpMTPSL2, MpCPT5 alone and MpMTPSL2 alone. Peak 1 and peak 2 are identified as α-phellandrene (peak 1) and D-limonene (peak 2), respectively.**

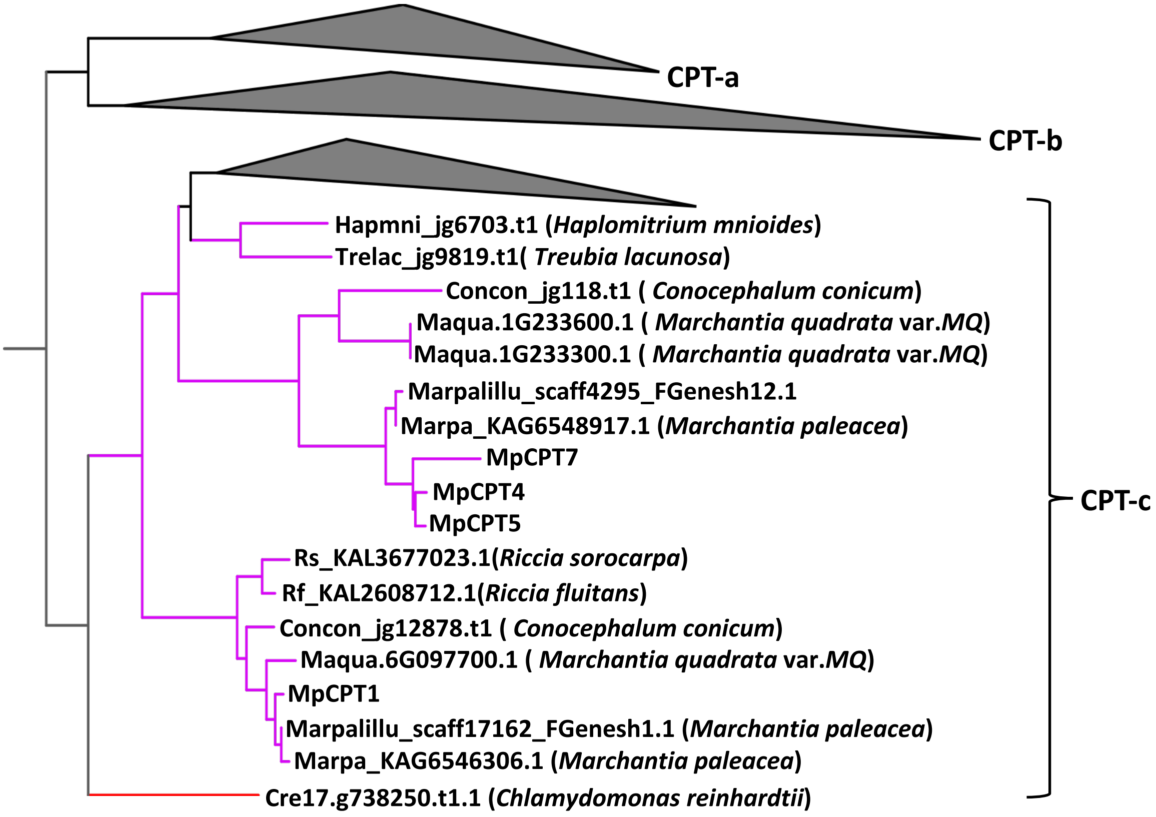

**Figure S13. Phylogenetic tree of CPTs.** The tree shown is the same as in Figure 7, with the CPT-a, CPT-b, and vascular plant CPT clades collapsed. Only CPT-c members from liverworts are shown.

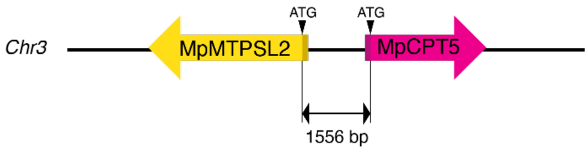

**Figure S14. Chromosomal organization of *MpMTPSL2* and *MpCPT5*.** Both genes are localized on chromosome 3 of *Marchantia polymorpha*. These two genes are in opposite orientation. The intergenic region from the ATG start codon of MpMTPSL2 to that of MpCPT5 is 1,556 bp.

**Table S1.** ***CPT*** **genes in** ***M. polymorpha***

| Gene name | Phytzome ID | Marchantia.info ID |
|---|---|---|
| *MpCPT1* | Mapoly0012s0016 | Mp8g02190.1 |
| *MpCPT2* | Mapoly0023s0020 | Mp2g10510.1 |
| *MpCPT3* | Mapoly0087s0025 | Mp4g05660.1 |
| *MpCPT4* | Mapoly0142s0037 | Mp3g18560.1 |
| *MpCPT5* | Mapoly0142s0042 | Mp3g18510.1 |
| *MpCPT6* | Mapoly0025s0048 | Mp2g26360.1 |
| *MpCPT7* | Mapoly0121s0026 | Mp3g23980.1 |
| *MpCPT8* | Mapoly0014s0198 | Mp1g10280.1 |

**Table S2. List of plants analyzed for the CPT family**

| Lineage | Species |
|---|---|
| Angiosperm | *Arabidopsis thaliana* |
| | *Solanum lycopersicum* |
| | *Oryza sativa* |
| | *Sorghum bicolor* |
| | *Amborella trichopoda var. SantaCruz_75* |
| Gymnosperm | *Thuja plicata* |
| Fern | *Adiantum capillus-veneris* |
| | *Alsophila spinulosa* |
| | *Azolla caroliniana* |
| | *Marsilea vestita* |
| | *Salvinia cucullata* |
| Lycophyte | *Diphasiastrum complanatum* |
| | *Selaginella moellendorffii* |
| | *Selaginella kraussiana* |
| Moss | *Funaria hygrometrica* |
| | *Physcomitrium patens* |
| | *Pohlia nutans* |
| | *Sphagnum fallax* |
| | *Sphagnum magellanicum* |
| | *Takakia lepidozioides* |
| Hornwort | *Anthoceros agrestis [Oxford]* |
| Liverwort | *Marchantia polymorpha* |
| | *Marchantia quadrata var. MQ* |
| | *Marchantia paleacea* |
| | *Conocephalum conicum* |
| | *Haplomitrium mnioides* |
| | *Treubia lacunosa* |
| | *Riccia fluitans* |
| | *Riccia sorocarpa* |
| Green algae | *Chlamydomonas reinhardtii* |

### Table S3. Primers used in this study.

| Primer name | Sequence (5' to 3') | |
|---|---|---|
| **For protein expression constructs** | | |
| ProexMpCPT5FP | GGATCCATGGTTTCACTCTCCAGAACACTCGG | pET-32a |
| ProexMpCPT5RP | AAGCTTTCAGAACTTGGTCTGTCTTAGACC | |
| ProexMpMTPSL2FP | ATGGCGGCAAAAGCTCACAGGAGTTTC | pEXP5-CT/TOPO |
| ProexMpMTPSL2RP | TTATCTCAAGTAGTGGTCCTCTTTTTG | |
| **For CRISPR/Cas9 construct** | | |
| MpCPT5.gRNA1_F | GCACCCAGCCTCTCGTTTCCTGGTGCGCCAGTGAAGTTTTAGAGCTAGAA | CRISPR/Cas9 gRNA |
| MpCPT5.gRNA1_R | TTCTAGCTCTAAAACTTCACTGGCGCACCAGGAAACGAGAGGCTGGGTGC | CRISPR/Cas9 gRNA |
| MpCPT5_PT_KO FW1 | TGCTTCATGGAGTCTGTGGTG | genome PCR |
| MpCTP5_PT_KO RV1 | TCTTCGTCTGGCTCAGATCTG | genome PCR |
| MpMTPSL2.gRNA1_F | GCACCCAGCCTCTCGGGAAGAACCCCGTATTTGGTGTTTTAGAGCTAGAA | CRISPR/Cas9 gRNA |
| MpMTPSL2.gRNA1_R | TTCTAGCTCTAAAACACCAAATACGGGGTTCTTCCCGAGAGGCTGGGTGC | CRISPR/Cas9 gRNA |
| MpMTPSL2_gRNA1-PCR-1L | TACCCGATTTCACTCATCTC | genome PCR |
| MpMTPSL2_gRNA1-PCR-4R | TCCGCAAATGCACTAAAGAC | genome PCR |
| **For subcellular localization** | | |
| LcMpCPT5FP | CACCATGGAGTCTGTGGTGGCAAGAT | Populus mesophyll protoplasts |
| LcMpCPT5RP | GAACTTGGTCTGTCTTAGACCAAG | |
| LcMpMTPSL2FP | CACCATGGCGGCAAAAGCTCACAGGAG | |
| LcMpMTPSL2RP | TCTCAAGTAGTGGTCCTCTTTTTGG | |
| **For transcriptional reporters** | | |
| proMpCPT5_fw | CTCAGGAGCGCGTATGTGAAGGGAAGTG | |
| proMpCPT5_rv | TAAGGTCTCGCATTGAAGCAGAGCCGACTGCTA AC | |
